# Designing Topographically Textured Microparticles for Induction and Modulation of Osteogenesis in Mesenchymal Stem Cell Engineering

**DOI:** 10.1101/2020.03.22.002279

**Authors:** Mahetab H. Amer, Marta Alvarez-Paino, Jane McLaren, Francesco Pappalardo, Sara Trujillo, Jing Qian Wong, Sumana Shrestha, Salah Abdelrazig, Lee A. Stevens, Jong Bong Lee, Dong-Hyun Kim, Cristina González-García, David Needham, Manuel Salmerón-Sánchez, Kevin M. Shakesheff, Morgan R. Alexander, Cameron Alexander, Felicity RAJ Rose

## Abstract

Mesenchymal stem cells have been the focus of intense research in bone development and regeneration. We demonstrate the potential of microparticles as modulating moieties of osteogenic response by utilizing their architectural features. Topographically textured microparticles of varying microscale features were produced by exploiting phase-separation of a readily-soluble sacrificial component from polylactic acid. The influence of varying topographical features on primary human mesenchymal stem cell attachment, proliferation and markers of osteogenesis was investigated. In the absence of osteoinductive supplements, cells cultured on textured microparticles exhibited notably increased expression of osteogenic markers relative to conventional smooth microparticles. They also exhibited varying morphological, attachment and proliferation responses. Significantly altered gene expression and metabolic profiles were observed, with varying histological characteristics *in vivo*. This study highlights how tailoring topographical design offers cell-instructive 3D microenvironments which allow manipulation of stem cell fate by eliciting the desired downstream response without use of exogenous osteoinductive factors.

## Introduction

Microparticles, also known as microcarriers, have gained significance as building blocks for tissue engineering strategies, in bioinks for three-dimensional (3D) bioprinting and for large-scale expansion of anchorage-dependent cells (*1*). Since they allow the formation of interconnected porous scaffolds and spatiotemporal release of bioactive factors, microparticles are an attractive tool for engineering complex tissues and biological interfaces. Understanding the correlation between microparticle-based cues and associated cellular responses is critical to achieve predictable outputs for translational applications. Tailoring surface properties of microparticles to direct differentiation in 3D provides the opportunity to transform cell delivery systems from passive mechanical supports to functional components of cell expansion processes and regenerative therapies.

Mesenchymal stem cells (MSCs), also known as multipotential stromal cells or mesenchymal stromal cells, are widely used in tissue engineering due to their multi-lineage potential and immunomodulatory effects (*2*). Since material properties influence MSC behaviour, tailoring the physical properties of microparticles makes it possible to achieve the desired biological responses. Topographical properties of the substrate play a key role in determining cellular responses (*3*), which have been shown to sometimes be more dominant than biochemical cues and stiffness (*4*). Therefore, the aim of this study was to explore the utilization of 3D architectural features of microparticles (i.e. topographical cues) for osteoinduction as a potential alternative for biochemical factors. We demonstrate the potential of capitalizing on cell-substrate interactions to develop cell-instructive microparticles, whereby tailored microparticle design can allow control over stem cell fate. This approach will facilitate the application of the ‘understand-predict-control’ engineering design principle within stem cell biology in a 3D platform, leading to improved microscale culture protocols for drug development and therapeutic applications. We have previously reported strategies to modify microparticle surfaces to improve cell adhesion (*5*), but the use of microcarriers with controlled topographies to drive stem cell differentiation has not been investigated. To test the capability of topographical designs on microparticles to induce osteogenesis, a range of viability and osteo-specific gene, protein and mineralization assays were performed using primary human mesenchymal stem cell (hMSCs) *in vitro*, and within non-healing murine radial bone defects *in vivo*.

## Results

### Fabrication of Textured Microparticles with Controlled Topographical Surfaces

Fabrication of microparticles using emulsion polymerization methods is amenable to scale-up for translational applications (*1*). Therefore, topographically textured polymer microparticles were fabricated using a drug-induced phase separation emulsion-solvent extraction method to investigate their ability to modulate MSCs differentiation *in vitro* and *in vivo*. Textured microparticles were produced by exploiting phase separation of fusidic acid (FA) from polymers during loss of solvent from an oil-in-water emulsion. Fusidic acid then dissolves leaving textured surfaces (**Fig. 1A, Supplementary Fig. 1C**). It was possible to design polymer-based microparticle systems with tunable surface features by changing polymer and drug composition. Towards the aim of producing optimized microparticles for bone regenerative engineering, polylactic acid (PLA) and poly(lactic-co-glycolic acid) (PLGA), two biocompatible polymers with varying biodegradability properties and suitability for implantation (*6*), were explored. By varying emulsion settings, microparticles with conventional smooth surfaces were produced, in addition to microparticles of two morphologies: dimpled and angular (**Fig. 1B**). Our studies have demonstrated that both microparticle and dimple sizes were dependent on polymer concentration and polymer/FA ratio (**Fig. 1C-D, Supplementary Table 1 and Supplementary Fig. 1B**). An increase in FA content slightly increased dimple size without altering particle size (**Supplementary Fig. 1A,B,D**). Dimpled microparticles were fabricated with different polymer/FA ratios and total concentrations in the organic phase (**Fig 1C and Supplementary Fig 1**). Sectioning of the fabricated microparticles revealed a largely non-porous internal structure, with a small number of sub-micron sized pores (**Supplementary Fig. 1E**). Although PLGA-based microparticles displayed uniform dimpled morphologies, degradation was rapid, and the dimpled morphology was not preserved after days in culture media (**Supplementary Fig. 1F**). Therefore, PLA-based microparticles were chosen for further cell studies (**Fig. 1B and Table 1**).

**Figure 1.**
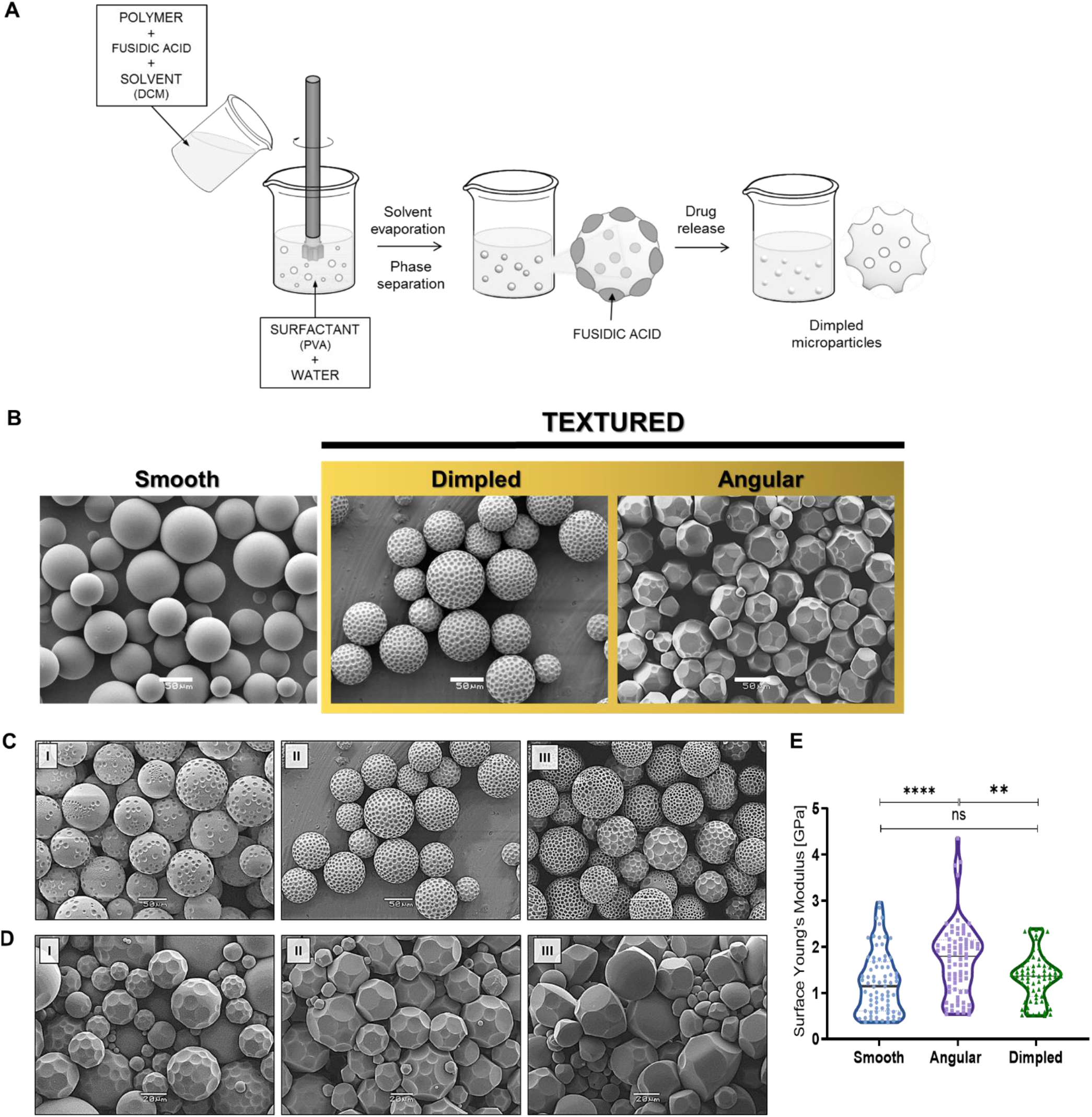
Fabrication and characterization of topographically textured microparticles. (A) Schematic representation of topographical microparticle fabrication by drug-induced phase separation oil-in-water solvent evaporation emulsion method (PVA: Polyvinyl acetate; DCM: Dichloromethane). (B) Representative SEM images of conventional smooth and the two topographically textured microparticle designs used in this study (Scale bar: 50 μm). (C) SEM images of dimpled PLA microparticles fabricated at I) 80/20, II) 70/30 and III) 60/40 PLA/FA ratios. PLA/FA concentration in the organic phase 10% (stirring speed 600 rpm; scale bar: 50 μm). (D) SEM images of angular PLA microparticles fabricated at I) 90/10, II) 75/25, III) 60/40 and IV) 40/60 PLA/FA ratios. PLA/FA total concentration in the organic phase is 15% (stirring speed 1600 rpm; scale bar: 20 μm). (E) Violin plot showing the AFM indentation-based surface elastic moduli measurements on loose microparticles (One-way ANOVA using Tukey’s *post-hoc* test; \*\**p*=0.0019, \*\*\*\**p*<0.0001)

**Table 1.** Properties of topographical microparticles used for assessment of cellular responses

|  |  | SMOOTH | DIMPLED | ANGULAR |
| --- | --- | --- | --- | --- |
| EMULSION<br>SETTINGS | FA/PLA ratio | 0/100 | 30/70 | 43/57 |
|  | RPM | 2000 | 600 | 1800 |
|  | Polymer concentration (%) | 20 | 10 | 20 |
| PARTICLE<br>PROPERTIES | Particle size (µm) | 58.2 ± 15.3 | 58.5 ± 13.2 | 43.4 ± 15.4 |
|  | Dimple size (µm) | - | 5.2 ± 2.2 | 12.0 ± 3.9 |
|  | Nº dimples/particle | 0 | 100 - 300 | 20 - 40 |
|  | BET surface area (m<sup>2</sup>/g) | 0.0486 ± 0.0022 | 0.1002± 0.0017 | 0.0944 ± 0.0017 |

### Characterization of Microparticles

Smooth, angular and dimpled particles displayed water contact angles of 87.96 ± 4.58, 92.25 ± 4.30 and 90.69 ± 2.89° respectively, consistent with previous reports on PLA (*7*). Surface areas (m^2^/g) of microparticles (**Table 1**) were calculated from the adsorption branch of Krypton isotherms at −196°C using the Brunauer-Emmett-Teller (BET) model.

Atomic force microscopy (AFM) was used to assess surface elasticity and roughness of microparticles. **Fig. 1E** shows that mean Young’s moduli were significantly higher for angular microparticles (1.72 ± 0.79 GPa) relative to smooth and dimpled microparticles (1.22 ± 0.67 and 1.33 ± 0.51 GPa, respectively). AFM measurements revealed elastic moduli values similar to those reported for collagen type I fibers (*8*) and bone (*9*). There were no significant differences between designs when assessed as overall mean values of the independent experimental repeats. To allow the correlation of data with observed cellular responses, measurements were also carried out on microparticles sintered into a disc, which was subsequently used for cell studies (**Fig. 2**). There were no significant differences between Young’s moduli of all different microparticle types when assessed within the disc format used for cell studies, nor between loose microparticles and their corresponding sintered disc format (**Fig. 2D**). Additionally, there were no significant differences in Young’s moduli after incubation with cell culture media (**Fig. 2D**). **Supplementary Fig. 2A** presents AFM topography images and 3D roughness profiles of each of the microparticle designs. Smooth microparticles displayed the lowest average surface roughness (R_*a*_) values while angular microparticles presented the highest ones. Interestingly, data revealed large differences in R_*a*_ values within the dimples of the textured dimpled particles compared to the exterior surface around the dimples (**Table 2**).

**Figure 2.**
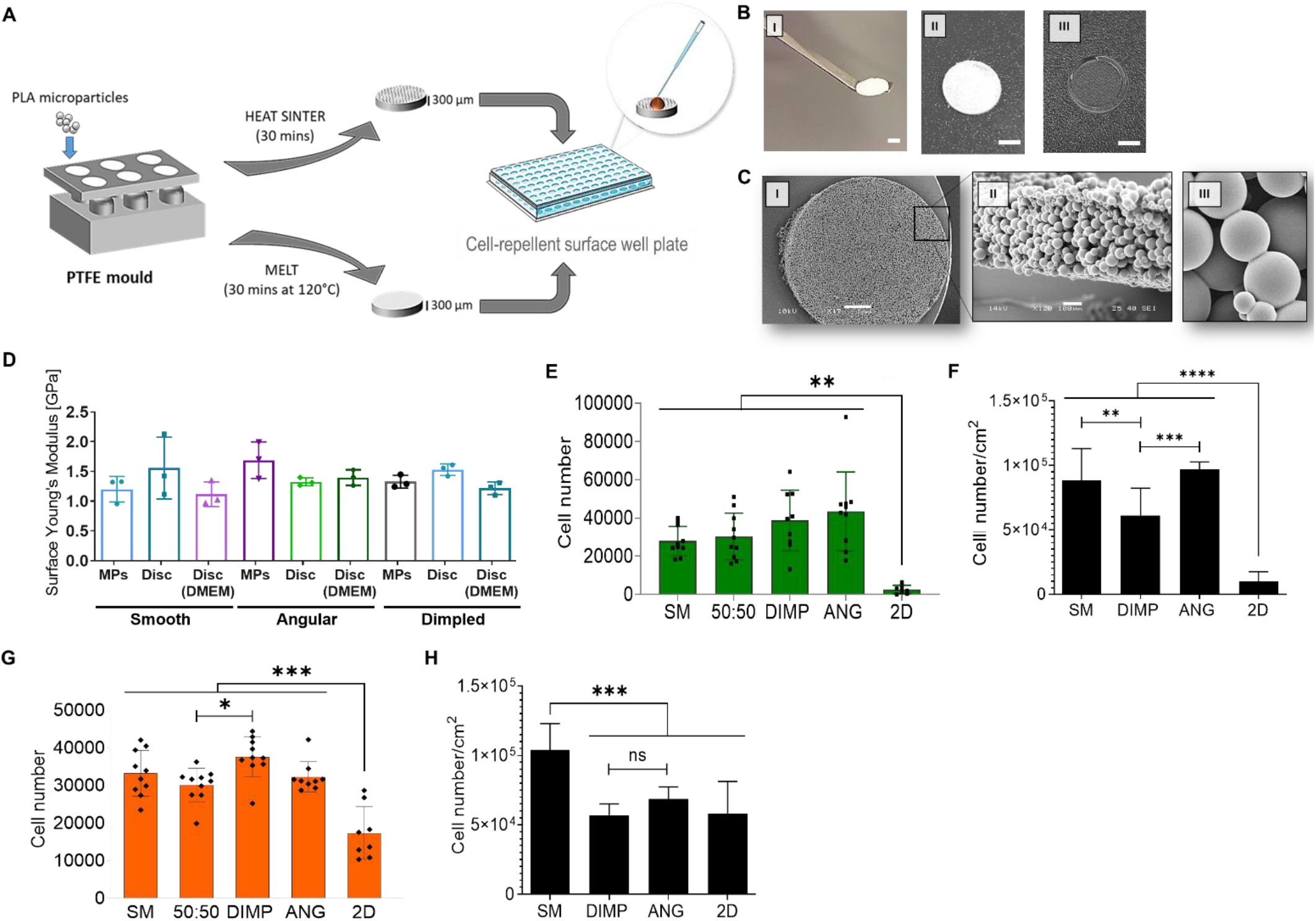
Cell attachment on the smooth and topographically textured microparticles using a planar cell culture platform. (A) Schematic showing the method used to prepare a planar presentation of the particles (‘2.5D’) for use in cell studies. (B) Sintered microparticle-containing discs (I, II) were 6.2 mm in diameter and approximately 300 μm-thin (Scale bar = 2mm). Discs made from melted microparticles were used as planar 2D controls (III). (C) SEM images showing top (I; scale = 1mm) and side (II; scale = 100 μm) views of the 2.5D planar discs used to assess cell response. A close-up of the sintered microparticles is shown (III). (D) Surface Young’s moduli measurements for loose microparticles versus sintered disc formats, measured dry and in DMEM culture media. (E) Attachment of primary human mesenchymal stem cells (hMSCs) from 3 different donors in 3 independent experiments, assessed using CyQuant, on various 2.5D disc topographies four hours post-seeding (*n* ≥ 8; one-way ANOVA and Tukey’s *post-hoc* test; *p*≤0.0044). (F) Cell attachment data corrected for surface areas to give cell numbers/cm^2^ (one-way ANOVA and Dunnet’s *post-hoc* test versus SM; \**p*=0.012, \*\**p*=0.006, \*\*\*\**p*<0.0001). (G) Attachment of primary hMSCs on the various 2.5D disc topographies four hours post-seeding, quantified using PrestoBlue (four independent experiments; 2 donors; *n* ≥ 8; one-way ANOVA and Tukey’ *post-hoc* test; \**p*=0.027, \*\*\**p*<0.0001). (H) PrestoBlue data corrected for surface areas (one-way ANOVA and Tukey’s *post-hoc* test; \*\**p*=0.002, \*\*\**p*=0.0006, \*\*\*\**p*<0.0001). SM: Smooth microparticle discs; DIMP: Dimpled microparticle discs; 50:50: Equal mixture of smooth and dimpled microparticles in discs; ANG: Angular microparticle discs; 2D: melted microparticle discs used as planar control.

**Table 2.** Values of surface roughness parameters (nm ± SD) of microparticles used in this study

| | $R_a$ | $R_q$ | $R_z$ |
| --- | --- | --- | --- |
| Smooth | $6.13 \pm 2.7$ | $7.89 \pm 3.2$ | $11.76 \pm 5.42$ |
| Angular | $70.25 \pm 7.2$ | $83.49 \pm 8.6$ | $102.5 \pm 10.2$ |
| Dimpled (outside dimples) | $9.16 \pm 1.6$ | $10.88 \pm 1.9$ | $13.55 \pm 2.9$ |
| Dimpled (within dimples) | $32.72 \pm 3.0$ | $37.84 \pm 3.1$ | $45.48 \pm 3.0$ |

*R<sub>a</sub>: Arithmetic mean roughness; R<sub>q</sub>: root mean square roughness; R<sub>z</sub>: average roughness between highest and lowest local* *minima*

To check for presence of residual fusidic acid (FA) content within the particles, release profiles of FA from textured particles was assessed at 37°C over 28 days following fabrication (**Supplementary Fig. 2B-C**). Results show initial burst release within the first hour, with 52% of FA released from the angular MPs and 22% from dimpled ones over 28 days. Since PLA does not degrade within this time span (**Supplementary Fig. 1F**), this explains the presence of residual FA entrapped within the polymer network. To limit any impact of residual fusidic acid on cell response, microparticles were washed with PBS under dynamic conditions for 7 days before use in cell studies. Furthermore, the influence of any possible residual FA was also assessed, as explained later.

To investigate the influence of microparticle topographical design on stem cell fate, a planar ‘2.5D’ presentation of the microparticles was developed (**Fig. 2A-C**). Microparticles were heat-sintered together into 300 μm-thin discs within a polytetrafluoroethylene (PTFE) mold, whilst maintaining their topographical characteristics. Corresponding flat ‘2D’ discs were produced by melting the microparticles at 120°C in the same manner to act as a planar control (**Fig. 2B**). One limitation of comparing different topographical designs is variable surface areas available for cell attachment. Since the weighing process led to fracture of these fragile discs, surface areas available for cell attachment were calculated using a mathematical approach. Modelling the microparticles as spheres within the disc (modelled as a cylinder) was not possible, as close packing of spheres (diameter σ) in a cylinder (diameter D) is a complex geometric question not possible to compute for D>2.873σ (*10*). Therefore, this packing problem was viewed as a 3D variant of the equal circle-packing problem in a bound container, where uniform microparticle packing and diameters were assumed (Mathematica *v* 11.3, Wolfram Research Inc.). Particle sizes and surface area measurements above (**Table 1**) were used to estimate the available (top hemi-spherical) surface areas available for cell seeding, which were calculated as 0.32, 0.66 and 0.47 cm^2^ for smooth, dimpled and angular discs respectively (calculations shown in **Supplementary Table 2**). For differentiation studies, effective cell seeding density was ≈5,000-10,000 cells/cm^2^. This range was used as the impact of variation in initial cell seeding density (cells/cm^2^) within the aforementioned range has been reported to have no influence on expression of osteocalcin and mineralization in MSCs (*11*–*13*).

### Varying Cell Adhesion Is Mediated by Different Integrins on Diverse Topographical Designs

When employing microparticles as cell carriers, cell adhesion is critical. DNA-based quantification of cell attachment on microparticles at 4 hours post-seeding was significantly higher than on planar 2D discs, with higher total cell attachment on dimpled microparticles relative to smooth ones (**Fig 2E**). Data showed significantly lower normalized attachment densities to dimpled microparticles and 2D planar controls relative to smooth microparticles (**Fig 2F**). Results were confirmed using the PrestoBlue assay, a metabolic activity-based measure of cell attachment (**Fig. 2G**), where normalized cell attachment was significantly lower on textured microparticles relative to smooth ones (**Fig. 2H**).

Cell morphology was influenced by different topographies, with cells spreading on smooth surfaces and adopting more rounded or elongated morphologies on textured microparticles (**Fig. 3A**). Differences observed in cell spreading was hypothesized to be due to preferential use of different integrins for adhesion to varying topographical designs. Therefore, cell adhesion was investigated further to identify which integrin(s) mediated cell-microparticle interactions. The ability of anti-integrin blocking antibodies to interfere with hMSCs adherence to smooth and dimpled microparticles, along with flat tissue culture-plastic (TCP) substrates, was studied. All tested anti-integrin antibodies blocked ≈75% of cell adhesion to smooth microparticles and ≈60% of adhesion to TCP substrates (**Fig. 3B**). In contrast, results suggested that hMSCs attachment to dimpled microparticles was mediated primarily via α_5_ and α_v_β_3_ only among integrins tested, with a significant difference in hMSCs attachment to smooth and dimpled microparticles with anti-α_2_ blocking antibodies (**Fig. 3B**). This demonstrates that binding to topographically textured particulate surfaces significantly alters integrin binding, and therefore impacts downstream signaling.

**Figure 3.**
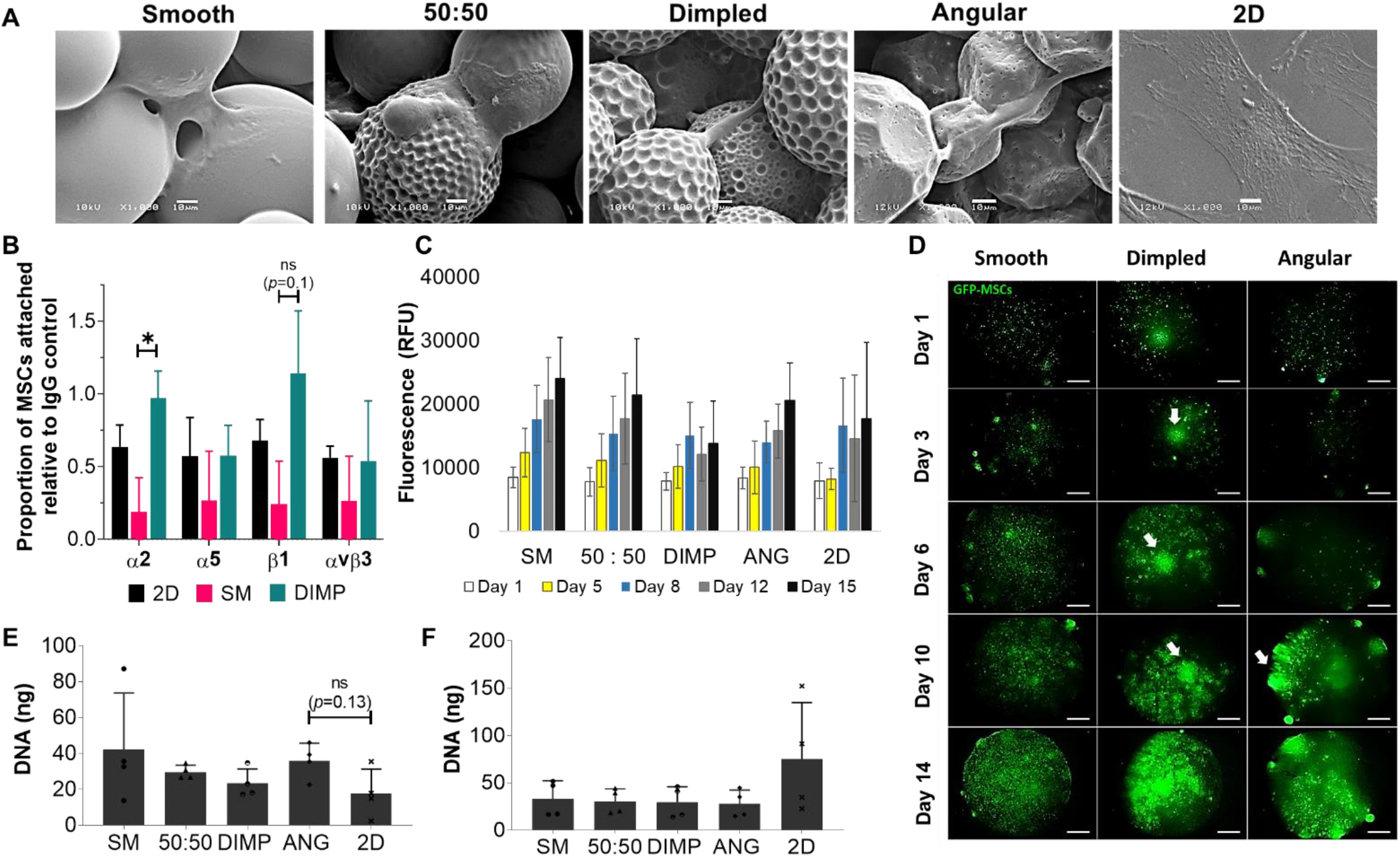
Adhesion and proliferation of hMSCs on varying microparticle designs. (A) SEM images showing morphology of primary hMSCs 4 hours post-seeding on the planar microparticle-containing discs (scale bar = 10 μm). (B) Adhesion of hMSCs to 2D TCP controls and 2.5D discs after pre-incubation with blocking antibodies against various integrins and incubated for 30 min. hMSCs pre-incubated with isotype IgG were used as controls (*n* = 4; 3 donors; 2 independent experiments; \**p*=0.03). (C) Proliferation of primary hMSCs, assessed using PrestoBlue on various 2.5D microparticle topographies (3 donors; *n* ≥ 6). (D) Fluorescence microscopy images displaying the proliferation of GFP-labelled hiMSCs on various 2.5D microparticle topographies (Scale bar = 1mm). Cell clusters are highlighted using white arrows. (E) DNA assessed at day 14 post-seeding, quantified using the Quant-iT PicoGreen dsDNA assay (2 donors, *n* = 4; two independent experiments; Kruskal-Wallis with Dunn’s *post-hoc* test). (F) DNA assessed at day 21 post-seeding, quantified using the Quant-iT PicoGreen dsDNA assay (*n* = 4, 2 donors). SM: Smooth microparticle discs; DIMP: Dimpled microparticle discs; 50:50: Equal mixture of smooth and dimpled microparticles in the discs; ANG: Angular microparticle discs; 2D: melted microparticle discs unless stated otherwise.

Cells proliferated on microparticles for 15 days, where PrestoBlue indicated consistently lower metabolic activity on dimpled microparticles at day 12 relative to day 8 (**Fig. 3C**). This was hypothesized to be due to either: (A) a decrease in cell number after day 8, which was not reflected in DNA-based quantification on day 14 (**Fig. 3E**) nor images of GFP-labelled hiMSCs on dimpled microparticles-containing discs (**Fig. 3D**); or (B) due to a decrease in cell metabolic activity. Cells were capable of increased proliferation at later time-points (**Fig. 3C**), signifying a transient metabolic decrease. Additionally, cell aggregation was observed on textured microparticles (**Fig. 3D**), while MSCs spread evenly over smooth microparticle discs. DNA-based quantification of cell numbers 14 days post-seeding showed similar cell numbers on all substrates, with marginally higher cell numbers on angular microparticles compared to 2D discs (*p*=0.1; **Fig. 3E**). There were no significant differences in cell numbers on microparticle-based substrates at day 21 (**Fig. 3F**) and were generally lower on microparticles than control, yet not statistically significant.

### MSCs on Textured Microparticles Express Varying Levels of Osteogenesis Markers in the Absence of Osteo-inductive Supplements

In the absence of exogenous osteo-inductive supplements, MSCs cultured on microparticle-based discs in basal culture media showed slightly higher levels of normalized ALP activity relative to 2D discs at day 7 (**Fig. 4A**). However, extracellular ALP levels secreted in the media were higher when cells were cultured on textured microparticles and 2D discs than on smooth microparticles (**Supplementary Fig. 3A**). Dimpled microparticles successfully induced osteogenic differentiation of hMSCs in basal media, as indicated by positive immunostaining (**Fig. 4B**) and increase in secreted levels of osteocalcin (OCN), an osteoblast-specific protein, after 2 and 3 weeks of culture in basal medium respectively (**Fig. 4C-D**). It is noteworthy that smooth microparticles showed slightly higher expression of osteocalcin than flat 2D discs (**Fig. 4B,C**), which correlates with reports on the ability of curved surfaces to promote osteogenesis (*14*). Quantified levels of OCN secreted by hMSCs on dimpled microparticles in basal culture medium were significantly higher than levels secreted by those cultured on other samples, as evidenced by enzyme-linked immunosorbent assay (ELISA) analysis at two time-points (*p*<0.05; **Fig. 4C-D**). OCN was expressed by hMSCs in all samples at day 14 post-seeding when cultured in osteoinductive media (**Supplementary Fig. 3B**). Mineralization was notably higher on textured microparticles, with positive von Kossa staining on textured microparticle discs. Interestingly, markedly higher mineralization was observed on angular microparticles (**Fig. 4E-F**), despite lower levels of OCN expression.

**Figure 4.**
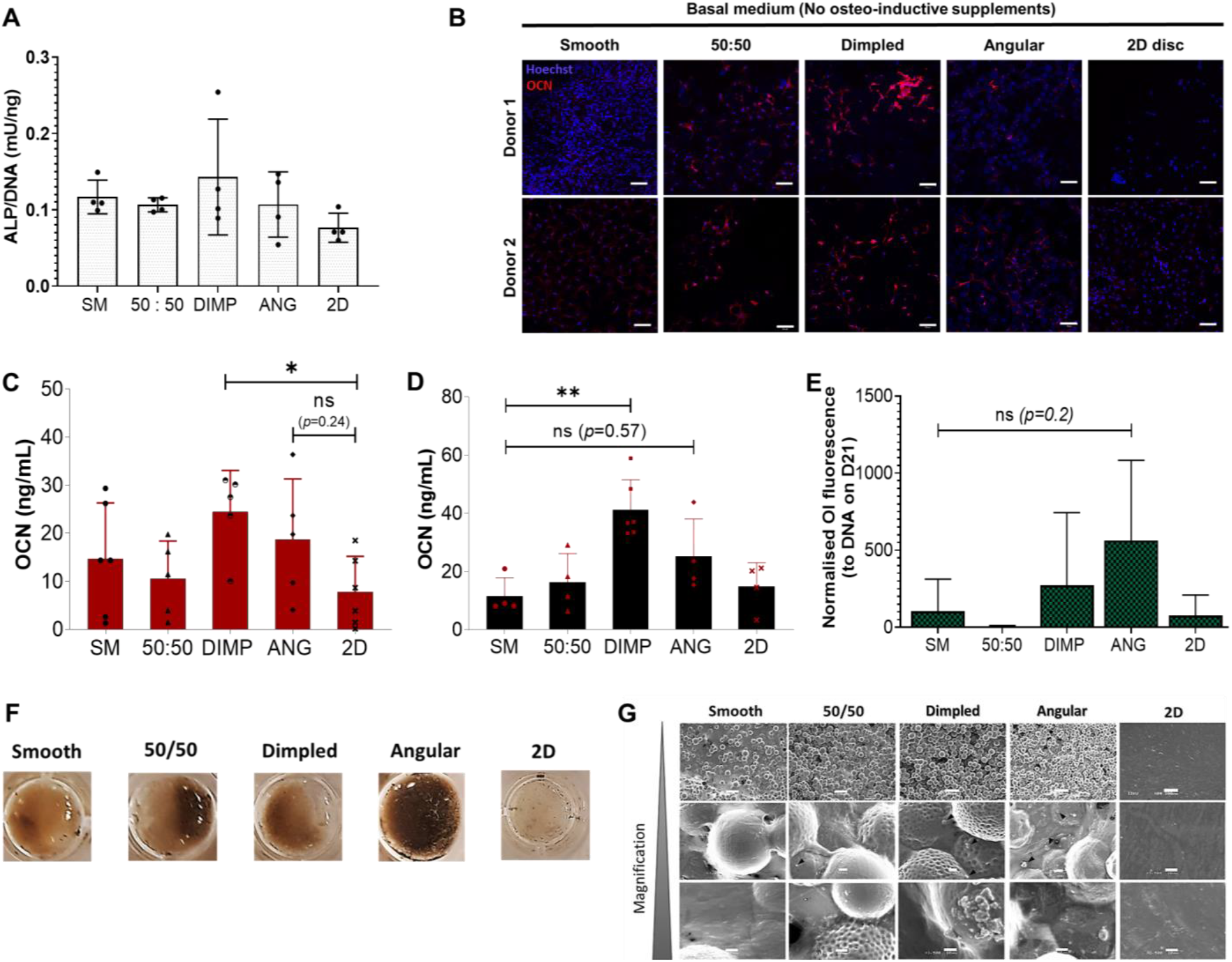
Textured microparticle surfaces drive the osteogenic differentiation of hMSCs. (A) ALP activity levels in cell lysates after normalizing to DNA content, of primary hMSCs cultured in basal culture media (no osteogenic supplements) for 7 days. Values shown are mean ± SD (*n* = 4 in two donors). (B) Maximum intensity projection images of z-stack sections obtained by immunostaining and confocal microscopy of primary hMSCs cultured on the various ‘2.5D’ discs in basal culture media for 14 days. Cell nuclei (Hoechst) are shown in blue and osteocalcin (OCN) in red. Microparticles, especially textured ones, were slightly auto-fluorescent in the blue channel (Scale bar = 100 μm). (C) Secretion of osteocalcin (OCN) in basal culture media of hMSCs, from two donors, cultured on the various discs at day 14 was detected using ELISA (*n* ≥ 5; two independent experiments; one-way ANOVA & Dunnett’s *post-hoc* test; \**p*=0.035). (D) Secretion of OCN in basal culture media of hMSCs, from two donors, cultured on the various discs at day 21 (*n* ≥ 4, two independent experiments; Kruskal-Wallis test & Dunn’s *post-hoc* test; \*\**p*=0.0088). (E) Graph showing average hydroxyapatite mineralisation on hMSCs-seeded discs, assessed using OsteoImage (OI) assay and normalized to DNA at day 21 (*n* = 4; 2 donors; Dunnett’s *post-hoc* test relative to SM). (F) Von Kossa staining of primary hMSCs-seeded discs, representative of two donors in 2 independent experiments. (G) Scanning electron microscopy images showing different areas of matrix deposition and nodule formation on hMSC-seeded discs after 21 days of culture in basal culture medium. Scale bars represent 200 μm (first row) and 10 μm (2^nd^ and 3^rd^ rows). Black arrowheads indicate nodule-like deposits. (SM: Smooth microparticle discs; DIMP: Dimpled microparticle discs; 50:50: Equal mixture of smooth and dimpled microparticles in discs; ANG: Angular microparticle discs; 2D: melted microparticle-based flat discs).

Genetic profiling supported these results, with differential gene expression between hMSCs cultured on TCP, smooth and dimpled microparticles identified. Genes for *COL10A1* were differentially upregulated in dimpled microparticle samples relative to TCP (5.5 fold; FDR=0.064), together with *BGN* (4.4 fold; FDR=0.048) and *transforming growth factor β receptor 2* (*TGFBR2;* 1.71 fold; FDR=0.031) when cultured on smooth microparticles (**Fig. 5A and Supplementary Fig. 4A**). Cells cultured on smooth microparticles displayed an upregulation of BGN. *TGFBR2*, encoding for transforming growth factor-beta (TGF-β) receptor type 2, was upregulated in cells cultured on smooth and dimpled microparticles. Interestingly, a change in expression of several osteogenesis-related genes were notably different on dimpled versus smooth microparticles (**Fig. 5A**). Altered genes included *COL4A3* (5.23 fold in smooth versus 1.21 fold in dimpled), EGF (0.41 fold change in dimpled versus 1.14 fold in smooth), and GDF10 (0.6 in dimpled versus 1.84 in smooth). Interestingly, *BMP-6* expression was downregulated in cells cultured on dimpled samples only (0.56 fold in dimpled versus 1.01 in smooth; **Fig. 5A**), despite being reported to be a potent osteogenesis inducer (*15*). Using hierarchical clustering with Euclidian distance measure, distinct clustering was observed containing either TCP, smooth microparticle or dimpled microparticle samples only (**Supplementary Fig. 4B**). Together, these results demonstrated that hMSCs differentiated towards an osteogenic lineage on textured microparticles, with varying levels of osteocalcin expression and mineralization.

**Figure 5.**
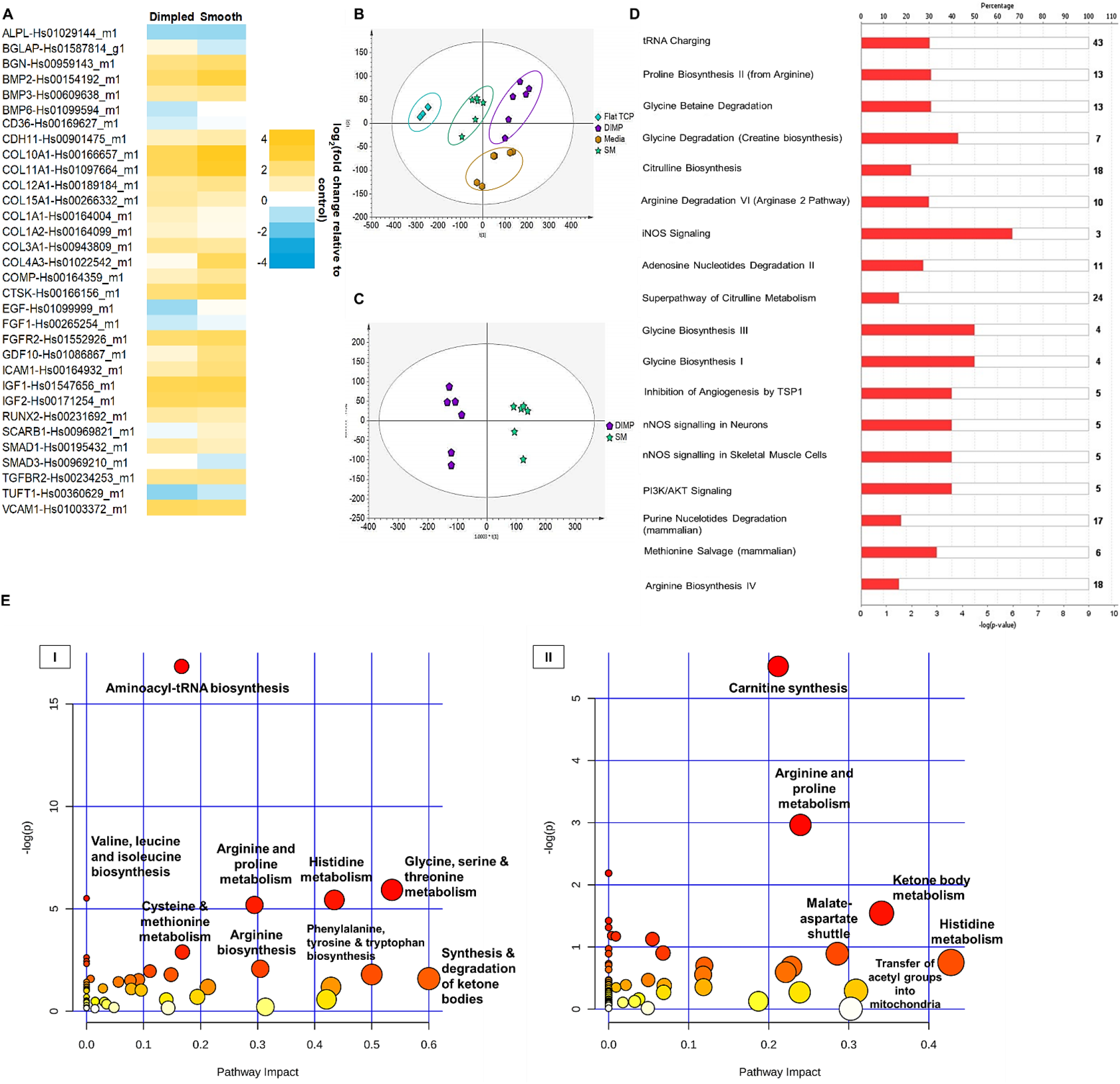
Genetic and metabolic profiles of hMSCs are significantly altered when cultured on dimpled microparticles. (A) Gene profiling revealed up- and down-regulated genes in hMSCs cultured on smooth and dimpled microparticles relative to control (cultured on TCP). Relative expression for selected genes is plotted as log2 (fold change) relative control after normalization against housekeeping genes. Yellow indicates upregulated and blue indicates downregulated genes (*n* = 2; 2 independent experiments). (B) PCA scores plot showing clear separation between metabolome profiles (*n*=6 microparticles and media samples; *n*=4 TCP samples; 2 donors; 2 independent experiments). (C) OPLS-DA scores plot based on metabolic profiling of culture media of smooth (SM) versus dimpled (DIMP) microparticles two weeks post-seeding (R^2^=0.98; Q^2^=0.97). (D) Top canonical pathways affected by culturing on dimpled versus smooth microparticles, generated using Ingenuity Pathway Analysis (IPA), as predicted by the differentially expressed metabolites (smooth *vs*. dimpled samples). Red color indicates upregulation. (E) Metabolic pathway analysis of the altered metabolites (smooth *vs*. dimpled) using MetaboAnalyst based on the KEGG (I) and SMPDB (II) databases. Color and size of each circle represents *p* values and pathway impact values respectively (Darker colors signify more influential changes in the pathway, and size represents the centrality of involved metabolites).

### Fusidic Acid Is Not Responsible for Osteoinductive Effects of Dimpled Microparticles

In order to delineate potential effects of any residual fusidic acid present after the fabrication process from the influence of topographical cues on stem cell fate, dimpled microparticle-conditioned media was employed to explore its effects on hMSCs differentiation. Two cell seeding densities were investigated to account for variability in initial cell attachment on different microparticle surfaces. As displayed in **Supplementary Fig. 3C-D**, there was no influence of the microparticles-conditioned media from smooth and dimpled microparticles on OCN expression or mineralization at days 14 and 21, respectively.

### Culturing hMSCs On Topographically Textured Microparticles Significantly Alters Their Metabolic Profile

Alterations in the genomic and proteomic landscapes are accompanied by changes in metabolic pathways, making metabolomics an appropriate method to assess cellular phenotype. Extracellular metabolite levels can be linked with intracellular metabolism (*16*). Samples of culture media were analyzed by liquid chromatography (LC)-mass spectrometry (MS) at day 15 post-culture to investigate adaptive metabolic changes occurring in hMSCs cultured on various microparticle designs and 2D. Data was analyzed using principal component analysis (PCA) and orthogonal partial least-squares discriminant analysis (OPLS-DA) to determine overall biological variation and identify differences in metabolic response. To avoid limitations associated with normalization to protein content, normalization was carried out to DNA quantification at the same time-point using IDEOM (*17*). PCA scores plot displays reproducible clustering of biological replicates for each group, and highlighted distinct differences in metabolic profiles of hMSCs cultured on different substrates (**Fig. 5B**). Unconditioned fresh media samples showed distinct clustering from samples in the presence of hMSCs (**Supplementary Fig. 4C**). Models were evaluated using cross-validation and goodness of fit (R^2^) and predictive ability (Q^2^) (0.98 and 0.97, respectively), indicating successful establishment of reliable models. PCA showed that exposure to either smooth or textured microparticles resulted in cells transitioning to a metabolic state different from that of TCP-cultured cells (**Fig. 5B**). Additionally, dimpled microparticle samples were clearly separated from smooth microparticle samples on the OPLS-DA scores plot, indicating altered metabolic states of hMSCs cultured on dimpled microparticles relative to smooth ones (**Fig. 5C**). Both microparticle types were also significantly separated from TCP-cultured cells on OPLS-DA scores plots (**Supplementary Fig. 4D**). Mass ions with VIP scores >1.0 were considered key differential metabolites, which was combined with Students *t*-test with false discovery rate correction to assess statistical significance.

Metabolites with significant differences between smooth and dimpled groups were identified as potential biomarkers of the influence of surface topographical design of microparticles. Metabolites including stearic acid, oleic acid, L-alanine and creatinine (**Supplementary Table 3**) were significantly higher in culture media of hMSCs on dimpled compared with smooth microparticle samples and controls. Two metabolites were found at remarkably higher concentrations in the media from cells cultured on textured microparticle samples relative to smooth microparticles. These were a trihydroxy epi-vitamin D3 derivative (194.8 and 124.1 fold higher in normalized and non-normalized data of dimpled samples relative to smooth, respectively, *p*<0.0001) and PG (20:1(11Z)/0:0), a glycerophospholipid (LPG(20:1); 70.6 and 45.0 fold higher in normalized and non-normalized data of dimpled compared to smooth, respectively; *p*<0.0001). Palmitic, stearic and oleic acids are major fatty acid components in human plasma lysophosphatidic acid (LPA)(*18*), which were also revealed to be significantly increased in the metabolic profile of dimpled media samples relative to smooth (**Supplementary Table 3**). ERK, interleukin-1 (IL-1) and JNK pathways were found to be central within the major signaling networks identified by Ingenuity Pathway Analysis (**Supplementary Fig. 5**).

As anticipated during active differentiation (*19*), key pathways such as aminoacyl tRNA biosynthesis, amino acid metabolism and energy-based pathways were significantly upregulated overall in hMSCs cultured on dimpled compared with smooth microparticles (**Fig 5D,E**). Ingenuity pathway analysis (IPA) of canonical signalling suggested the osteo-inductive effect of dimpled microparticles may occur via enhanced activity of iNOS (**Fig. 5D**). Metabolic pathways of the identified differentially abundant metabolites between smooth and dimpled microparticles media samples were also examined using MetaboAnalyst. Pathway significance was determined from pathway enrichment analysis, and impact values were determined based on the influence of metabolite(s) on a pathway’s function. A significantly affected pathway was arginine and proline metabolism, including L-glutamate-5-semialdehyde and L-citrulline. Top upstream regulators, identified by IPA based on expected effects between transcriptional regulators and known target genes in the Ingenuity^®^ database, included *ZC3H10*, consistent with its activated state.

### Textured Microparticle Designs Induce Distinct Bone Regeneration Patterns in Vivo

To evaluate the bone-forming potential of topographically-patterned microparticles *in vivo*, a murine non-healing radial bone defect model was used(*20*). Polyimide sleeves, with pores along the sides, were filled with either smooth, dimpled or angular microparticles in a collagen gel carrier (**Fig. 6A**). Low elastic modulus collagen type I was used as a biocompatible carrier system. As the aim was to evaluate the osteogenic reconstruction potential of microparticle design, eight weeks were deemed adequate to interpret differences based on histology.

**Figure 6.**
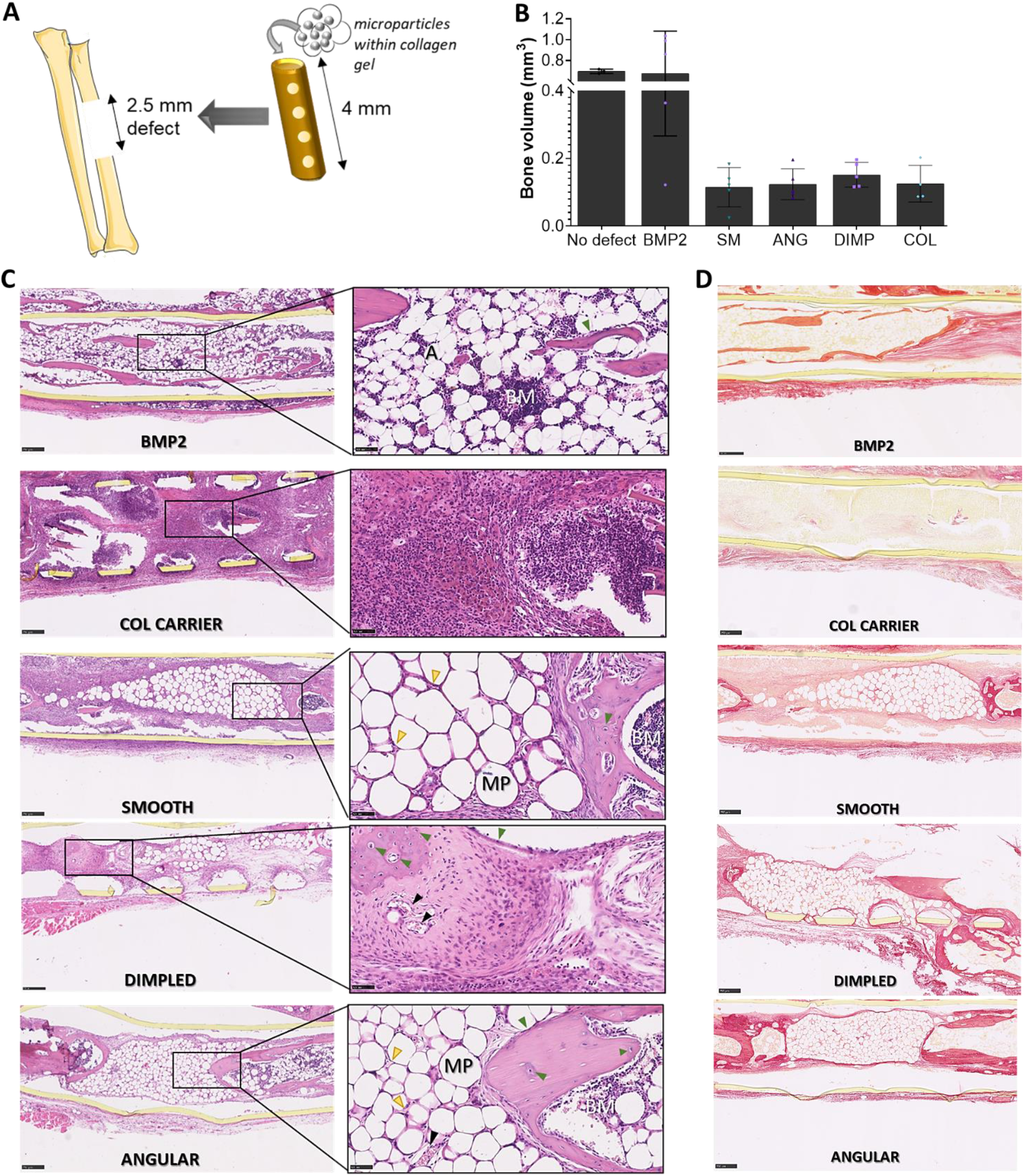
Influence of microparticle design on bone regeneration in a non-healing defect without exogenous cells or growth factors. (A) Samples were placed into cylindrical, porous polyimide sleeves and implanted in a non-healing defect (2.5 mm) in a murine radius. Implant tubes were filled with either 3 μL of microparticles suspension in collagen, 75 μg/mL BMP-2 in collagen or plain collagen (*n* = 5 per condition). (B) Measures of bone volume within defects, assessed using μCT, after 8 weeks (SM: smooth, DIMP: dimpled, ANG: angular, BMP-2 as positive control, COL: collagen carrier only). (C) Histological images showing sections of 8-week radial samples stained with H&E staining (scale bar: 250 μm), with areas of interest magnified (scale bar: 50 μm). Green arrowheads highlight bone elements (osteoblasts, osteocytes), yellow arrowheads show cell infiltration in between microparticles packed within the defect, black arrowheads indicate formed blood vessels within the defect (BM: bone marrow elements; MP: space occupied by microparticle; A: adipocytes). (D) Sections of 8-week radial samples stained with PicroSirius Red, for detection of the collagen matrix network in dark red (Scale bar: 250 μm).

We examined: (i) smooth, (ii) dimpled, (iii) angular microparticles, in addition to (iv) collagen gel carrier only, and (v) collagen gel carrier supplemented with 75 μg/mL bone morphogenetic protein-2 (BMP-2). We evaluated bone formation by microcomputed tomography (μCT) and histological examination at eight weeks post-implantation. Quantification of bone volume by μCT (**Fig. 6B and Supplementary Fig. 6A**) revealed that only BMP-2-loaded collagen resulted in significant repair and bridging of the defect. As expected, microparticles and collagen carrier alone did not promote the same level of bone growth as the BMP-2 impregnated collagen gel, although dimpled microparticles displayed a slightly higher mean bone volume relative to other designs (**Fig. 6B**).

Histological analysis at 8 weeks showed fibrous tissue with no significant bone formation within the defect for all the conditions except BMP-2, where both bone formation and bone marrow establishment in the center of the defect were noted, including hematopoietic elements (**labelled BM**; **Fig. 6C**) and adipocytes (**Fig. 6C**). Higher-magnification images within the defect confirmed different tissue characteristics among microparticles and the plain collagen carrier. With plain collagen, we observed mostly fibrous connective tissue with patches of inflammatory granulation tissue (**Fig. 6C**) and almost no collagen deposition in the ECM, as indicated by Picrosirius Red staining (**Fig. 6D**). Despite the presence of tightly packed microparticles within the implant tube, cell infiltration was clearly noted within the defects (**yellow arrowheads; Fig. 6C**). Some bone formation could be observed in defects containing smooth microparticles, possibly attributable to the stiffness of PLA stimulating osteogenesis to some degree (*4*). With dimpled and angular microparticle-containing implant tubes, cells were organized into larger endosteum-like structures as in the bone marrow, displaying osteoblasts and osteocytes in lacuna, and compact bone (**Fig. 6C and Supplementary Fig. 6C**). Abundant collagen deposition, indicated by Picrosirius Red staining, was noted in regions that exhibited new bone formation (**Fig. 6D**). However, there was an evident lack of hematopoietic bone marrow elements observed in the defects treated with dimpled microparticles compared to smooth and angular microparticles, despite the presence of supportive networks of newly-formed trabeculae for hematopoiesis to occur (**Fig. 6C**). Alcian Blue staining (**Supplementary Fig. 6B**) indicated the presence of sulfated glycosaminoglycans, which was especially notable within defects containing angular microparticles. Additionally, the presence of a high number of well-organized blood vessels within defects containing both types of textured microparticles was observed (**black arrowheads in Fig. 6C and Supplementary Fig. 6C**).

## Discussion

Research has focused on characterizing the interactions of MSCs with substrates of varying mechanical, surface chemistry, and topographical properties (*3*, *4*, *20*–*22*). On the other hand, less research has capitalized on how these matrix-based biophysical cues could be used to direct MSCs towards specific lineages for basic research and translational applications (*2*). Our results are important for formulating design criteria for microparticle-based environments to promote *ex vivo* control of cell fate for stem cell biomanufacturing and disease model development. In this work, we investigate the feasibility of exploiting cell-substrate interactions to develop cell-instructive platforms for driving osteogenic differentiation of hMSCs in 3D without the use of costly growth factors. We provide evidence herein of the *in vitro* topographically-patterned 3D niche microenvironments driving MSCs to go down an osteogenic lineage. Our findings indicate that MSC morphology, adhesion, proliferation, osteogenic lineage specification, and metabolic profile can be regulated using microengineered substrate-based cues in the form of microparticle topographical design.

When employing microparticles as cell carriers, cell adhesion is critical. Increased total cell attachment to microparticles compared to 2D controls is expected due to their increased surface area-to volume ratio. Significantly lower normalized attachment densities to dimpled microparticles relative to smooth microparticles agrees with a previous study describing anti-adhesive properties of pits between 3-10 μm (*23*), which is similar to dimple sizes described herein. Results were confirmed using a metabolic activity-based measure of cell attachment, where normalized cell attachment was significantly lower on textured microparticles relative to smooth ones. Metabolism has also been reported to be lower in cells exposed to topographies relative to control substrates (*24*).

Morphological differences in the cultured MSCs can be explained by how the surfaces regulate intracellular tension. On the smooth microparticles, the stiff, smooth surfaces will resist contractile forces exerted by MSCs to cluster integrins, which facilitate actin polymerization and act as anchoring points. However, direct physical interference of the micro-scale topographical patterns with the establishment and maturation of focal adhesion blocks cell spreading (*23*). Differences observed in cell morphology and spreading was also hypothesized to be due to the preferential use of different integrins for adhesion to varying topographical designs. Integrins initiate numerous downstream signaling pathways that regulate cellular processes, including differentiation (*22*). Results suggested that hMSCs attachment to dimpled microparticles was mediated primarily via α_5_ and α_v_β_3_ only among integrins tested, with a significant difference in hMSCs attachment to smooth and dimpled microparticles with anti-α_2_ blocking antibodies. This demonstrates that binding to topographically textured particulate surfaces significantly alters integrin binding, and therefore impacts downstream signaling. Both α_2_ and α_5_ integrins are mechanosensitive molecules which enhance osteogenic differentiation in response to different spherical spatial boundary conditions (*25*). Moreover, signaling through α_v_β_3_ integrin plays a significant role in promoting osteogenesis (*26*).

The transient metabolic decrease of hMSCs cultured on the dimpled microparticles may reflect osteoblastic differentiation and/or calcium internalization at that time point (*27*, *28*). The higher cell numbers on angular microparticles may be due to reduced physical tension experienced by cells on the flat regions available, which may have allowed cells to grow faster compared to cells on convex surfaces (*29*). Cell aggregation observed during proliferation on textured microparticles may be due to varying patterns of cell adhesion caused by somewhat variable dimple sizes. Some dimples may have been more conducive for induction of differentiation than others (*30*). This may have led to creation of heterogeneous microenvironments within the same sample as early as day 3. Since adhesion-based signaling contributes to MSC differentiation in monolayer cultures (*31*), data herein suggests that cellular mechanoreceptors can be exploited to induce osteogenesis in 3D systems. Osteogenic differentiation is enhanced on compact 3D structures, where cells display smaller cell areas and higher major axis length (*3*), similar to cell morphologies observed on dimpled microparticles. Additionally, MSCs cultured on disordered topographies display elevated levels of osteocalcin (*21*). The results reported herein demonstrate that hMSCs differentiated towards an osteogenic lineage on textured microparticles, with varying levels of osteocalcin expression and mineralization. In the absence of exogenous osteo-inductive supplements, dimpled microparticles successfully induced osteogenic differentiation of hMSCs in basal media, as indicated by the significant increase in levels of OCN expression after 2 and 3 weeks of culture. It is noteworthy that smooth microparticles showed slightly higher expression of osteocalcin than flat 2D discs, which correlates with reports on the ability of curved surfaces to promote osteogenesis (*14*). Mineralization was notably higher on textured microparticles. Interestingly, markedly higher mineralization was observed on angular microparticles, despite lower levels of OCN expression. This can be correlated to previous studies *in vivo*, where osteocalcin-null mice were reported to exhibit increased mineral-to-matrix ratio (*32*), while mice overexpressing osteocalcin showed normal mineralization (*33*).

Differential gene expression between hMSCs cultured on TCP, smooth and dimpled microparticles was identified. Genes for *COL10A1* were differentially upregulated in dimpled microparticle samples relative to TCP. COL10 is associated with endochondral ossification and directing bone mineralization (*34*). Cells cultured on smooth microparticles displayed an upregulation of BGN, which regulates collagen fibrillogenesis (*35*). *TGFBR2*, encoding for transforming growth factor-beta (TGF-β) receptor type 2, was upregulated in cells cultured on both smooth and dimpled microparticles, suggesting the possibility of stiff, curved surfaces priming cells to be more responsive to TGF-β regulation of osteogenesis. A change in expression of several osteogenesis-related genes were notably different on dimpled versus smooth microparticles. Altered genes included *COL4A3*, which encodes for a type IV collagen component making up epithelial and endothelial cell basement membranes (*36*). Downregulated genes in MSCs cultured on dimpled microparticles included EGF, which promotes *ex vivo* expansion of MSCs without triggering differentiation, maintaining its stemness (*37*), and GDF10, which plays an inhibitory role in osteoblast differentiation (*38*). Interestingly, *BMP-6* expression was also downregulated in cells cultured on dimpled samples only, which may be related to its recently reported role in the inhibition of glucose production (*39*).

Extracellular metabolite levels can be linked with intracellular metabolism (*16*). Metabolomics data analyses suggest that hMSCs are both a producer and target of vitamin D3 metabolites when cultured on dimpled particulate surfaces, which suggests a potential role of vitamin D metabolism in topographically induced differentiation. While some mammalian cells can make their own 1,25-dihydroxyvitamin D (1,25(OH)_2_D_3_) and respond to it (*40*), hMSCs express low levels of 25(OH)D_3_-1α-hydroxylase and vitamin D receptor (*41*). Nevertheless, hydroxy-derivatives of vitamin D3 have been shown to stimulate osteogenesis (*41*). PGE2 and TGF-β_1_ levels were shown to be regulated by 1α,25(OH)_2_D_3_ in osteoblast-like cells, with effects synergistic with increasing substrate microroughness and absent on smooth surfaces (*42*). LPG (20:1(11Z)/0:0), a monoacylglycerophosphoglycerol, was also identified as a significantly increased metabolite when hMSCs were cultured on dimpled microparticles relative to smooth ones. Osteoblastic differentiation requires lipidomic remodeling, driven by polyunsaturated lipids with long acyl chains (*43*). Palmitic, stearic and oleic acids are major fatty acid components in human plasma lysophosphatidic acid (LPA) (*18*), which were revealed to be significantly increased in the metabolic profile of dimpled media samples relative to smooth. LPAs regulate extracellular-signal-regulated kinase (ERK) pathways, which are central to cell differentiation (*44*). This agrees with the major signaling networks identified by Ingenuity Pathway Analysis (IPA), where ERK, interleukin-1 (IL-1) and JNK pathways are central. Supporting literature reports that integrin-related signaling has a central role in the modulation of stem cell phenotype on nano-topographical surfaces, acting through ERK1/2 and Jnk, with LPA being vital in driving osteogenic differentiation (*19*). IPA of canonical signalling suggested the osteo-inductive effect of dimpled microparticles may occur via enhanced activity of iNOS, which is expressed in bone marrow stromal cells and osteoblasts, and is involved in the stimulation of osteoblasts by NO production (*45*). Other significantly affected pathways included arginine, which plays an important role in bone healing through the production of nitric oxide (*46*), and proline, a crucial component of collagen (*47*). Carnitine has also been linked to osteogenic differentiation (*48*). Ketone body metabolism is a significant contributor to mammalian energy metabolism within extrahepatic tissues when glucose is not readily available (*49*). Top upstream regulators identified by IPA included *ZC3H10*, consistent with its activated state. *ZC3H10* is a poorly characterized RNA-binding protein which regulates mitochondrial biogenesis (*50*), which correlates with the importance of mitochondrial dynamics in osteogenesis (*51*). More studies are needed to reveal how it regulates osteogenesis. Further experiments on intracellular metabolites will confirm mechanisms associated with the metabolic adaptation of hMSCs to topographically textured surfaces.

To evaluate the bone-forming potential of topographically-patterned microparticles *in vivo*, a murine non-healing radial bone defect model was used (*20*). This was a demanding set-up, where we aimed to investigate the contribution of topographically textured microparticles to bone regeneration without incorporation of any exogenous variables, such as cells or growth factors. With textured microparticle-containing implant tubes, cells were organized into larger endosteum-like structures and abundant collagen deposition. However, there was an evident lack of hematopoietic bone marrow elements observed in the defects treated with dimpled microparticles compared to smooth and angular microparticles, despite the presence of supportive networks of newly formed trabeculae to support hematopoiesis. We hypothesize that incorporating low quantities of growth factors within the microparticles will promote synergistic effects of topographical design and microparticle-mediated growth factor presentation (*20*), which will be investigated in future studies. This confirms that the appropriate microparticle design can maximize bone regeneration potential in clinical applications and is vital to support optimal bone growth. This should be chosen based on its ability to support new bone formation, alongside promotion of hematopoiesis and bone marrow restoration.

## Conclusions

This study offers novel insights into the substantial impact of topographical patterning of microparticles on hMSCs response. We demonstrate herein how hMSCs morphology, adhesion, proliferation, osteogenic lineage specification, metabolic profile and *in vivo* histological characteristics can be modulated using microparticle-based 3D topographical design. Such micro-scale control represents a crucial advance in our ability to study and regulate stem cell–microenvironment interactions in 3D. Designing cell-instructive microparticles with defined topographies provides a promising platform for investigating how microenvironmental cues modulate the development of 3D-engineered bones and to create *in vitro* high-throughput drug screening array systems. Combining such platforms into next-generation organoid and niche bioengineering protocols will enable improved system control and increase the ability to probe the multivariable complexity of biological systems. Overlaying microparticle-based bioengineering techniques to pattern micro-tissues and develop ‘multiplexed’ organoids can improve fidelity of developmental models, acting as mechanical cue providers to emulate the native cellular microenvironment for the development of *in vitro* models for basic and translational stem cell research. Moreover, this study provides key insights into the design of instructive, injectable scaffolds for bone regenerative repair.

## Materials and Methods

### Fabrication of Smooth and Textured Microparticles

Smooth poly(D,L-lactic acid) (PLA) microparticles (Evonik Industries; Mn 47000 gmol^−1^, IV~ 0.5 dL g^−1^) were prepared by a solvent evaporation oil-in-water emulsion technique. The organic phase, containing 1g of PLA in 5 mL of dichloromethane ((20% *w/v;* DCM; ≥99.8, Fisher Scientific) was homogenized (Silverson Homogenizer L5M) at 2000 rpm for 5 minutes in the aqueous continuous phase (100 mL) containing 1% *w/v* of poly(vinyl acetate-co-alcohol) (PVA; MW 13-23 kDa, 98% hydrolyzed; Sigma-Aldrich) as stabilizer. The resulting emulsion was stirred continuously at 500 rpm at room temperature for at least 4 hours to allow for solvent evaporation. To remove residual PVA, microparticles were centrifuged at 4500 rpm for 5 min and subsequently washed with deionized (DI) water three times. Cell strainers with pore sizes of 40 and 100 μm were used to separate microparticles in this size range and the collected microparticles were freeze-dried for storage at −20°C.

To produce textured microparticles, fusidic acid (FA; 98%, Acros Organics or F0756-#SLBN8134V, Sigma-Aldrich) was incorporated in the organic phase as previously described(*52*), with 30 wt% in polymer for fabrication of dimpled MPs and 43 wt% for angular MPs (FA/PLA total content of 10% *w/v* in DCM for dimpled MPs and 20% *w/v* in DCM for angular MPs). The dispersed phase was stirred at 600 rpm into PVA solution (1% *w/v*). After solvent evaporation, microparticles were washed and collected. Textured microparticles were obtained after FA release under dynamic conditions in phosphate buffered saline (PBS; 0.1M, pH 7.4, 0.15M NaCl) at 37°C over 7 days.

### BET Surface Area Measurements Using Kr Isotherms

Kr Isotherms were acquired using a Micromeritics ASAP 2420 using Kr as the adsorbate at −196°C. Approximately 0.5 g of sample was weighed into a sample tube and degassed at 37 °C for 15 hours under high vacuum to remove moisture and other adsorbed gases. Isotherms were taken from 0.07-0.27 relative pressure and BET specific surfaces area were calculated in this range using Microactive Software v5.00.

### Heat-Sintered 2.5D Discs for Cell Culture Studies

Microparticles were packed into a PTFE mold and heated to 70°C (higher than the glass transition temperature of PLA) for 30 min. This induces microparticles to bond with neighboring particles, forming a ‘2.5D’ 300 μm-thick disc (6.2 mm diameter). Discs were washed in PBS and sterilized by UV irradiation for 20 min. The ‘50:50’ discs denote discs made up of 50% smooth and 50% dimpled microparticles.

### Hydrophobicity Measurements

Particles were suspended in a mixture of water and ethanol and left to settle on glass slides to form a monolayer. Once dry, MPs were slightly sintered at 70°C for 10 min. Contact angles were measured with a CAM 200 instrument (KSV Instruments, Finland) using the sessile drop method at 25°C. Contact angles were calculated using Young–Laplace curve fit by CAM 200 image analysis software, and resulting right and left contact angles were averaged. A minimum of five repeat measurements were made for each surface type.

### AFM Elastic Moduli Measurements

The MFP-3D Standalone Atomic Force Microscope (AFM) (Oxford Instruments, Asylum Research Inc., CA) and AFM probe RTESPA-150 (Bruker Nano Inc., CA; half-angle 21°; spring constant 1.8 N/m) were used to obtain force-displacement curves. Derjaguin-Muller-Toporov (DMT) model was employed for extrapolating Young’s moduli from the linear region of the retracting curve by applying ordinary least squares method. To prepare PLA microparticle samples for analysis, a thin layer of epoxy glue (Huntsman Advanced Materials, Switzerland) was applied on a glass slide and then evened out using another glass slide. A small amount of microparticles (or discs) were placed on the glue. After a few minutes, the slide was flipped to remove unattached microparticles and then left overnight for glue hardening. Samples were tested in air, water and culture media at RT for force measurement. Before each series of measurements, instrument sensitivity was calibrated with the unloading curve slope of an F-Z curve of glass alone, whilst the instrument’s thermal fluctuations were used to extract the spring constant of the cantilever. For each measurement, at least 50 F-Z curves were recorded. Acquired data was converted into .txt files containing Deflection and *Z*-sensor points by the subroutine provided in the associated MPF3D-AFM software. The .txt files were processed in Microsoft Excel by the DMT model to fit the retracting curve and calculate E.

### AFM Topographical Analysis

The Dimension FastScan Atomic Force Microscope (Bruker Nano Inc., CA) and AFM probe RTESPA-150 (Bruker Nano Inc.; half-angle 21°; spring constant 1.8 N/m) were used to explore surface topographies of microparticles for surface roughness evaluation. AFM indentation tests were conducted on the uppermost portion of each sample’s surface. From detected deflections, a surface topography map is generated. PeakForce QNM technique, integrated in the Bruker Nanoscope software, was employed for nano-mechanical mapping. Prior to analyses, laser and detector alignment was set automatically. Samples were tested in air for surface mapping, where 256×256 pixels surface maps were recorded at 4 or 7 μm surface area for a total of 3 images per sample. Images were processed via Gwyddion software(*53*) for tip deconvolution and image artefact correction.

### Fusidic Acid Release

Fusidic acid (FA) release was assessed by placing microparticles in 15 mL of PBS at 37°C. At various time points, samples were centrifuged and the supernatant collected, replacing it with fresh PBS. Filtered samples were analyzed for concentration of fusidic acid using an HPLC-UV system (Waters 600 Pump, Waters 717 Autosampler and Waters 2996 Photodiode Array Detector). The column and samples were maintained at temperature of 40°C and 6°C, respectively. The mobile phase comprised acetonitrile:10 mM ammonium acetate buffer (pH 4.1) at 60:40, with flow rate of 0.6 mL/min. The stationary phase was a Waters Atlantis C18 2.1 × 150 mm, 5 μm particle size column (Milford, MA, USA) protected by a SecurityGuard 2 × 4 mm, 3 μm particle size (Phenomenex, UK). Bexarotene (CAS: 153559-49-0, LC Labs, USA) was used as an internal standard and chromatograms were observed at 220 nm.

### Primary Human Mesenchymal Stem Cell Culture

Primary human bone marrow-derived mesenchymal stem cells (hMSCs) were obtained from Lonza (Germany) and cultured in basal mesenchymal stem cell growth medium (MSCGM) supplemented with 10% (*v/v*) fetal bovine serum, 2% (*w/v*) L-glutamine and 0.1% (*w/v*) Gentamicin-Amphotericin (#PT-3001; Lonza, Germany) with 5% CO2 in air at 37°C. Lot numbers of hMSC batches obtained were #0000351482 (male), #0000411107 (female), #0000491129 (female) and #0000422610 (male), cultured as individual patient stocks. Cells used were between the third and sixth passages. These cells were tested for their ability to differentiate into osteogenic, adipogenic and chondrogenic lineages, and for expression of surface markers recommended by the International Society for Cellular Therapy(*54*). Routine passaging and differentiation procedures were performed according to the relevant Lonza Poietics™ protocols.

### Assessment of Cell Attachment and Metabolic Activity

Cells were seeded on sintered discs in cell-repellent CellStar^®^ 96-well plates (Greiner Bio-One) at 3×10^4^ cells/well and incubated for 4 hours. Discs were then washed gently with PBS to remove non-adherent cells. DNA-based quantification of cell attachment was determined by CyQUANT NF Cell Proliferation Assay (ThermoFisher Scientific, UK) using a Tecan Infinite M200 microplate reader (Tecan, UK), with λ_exc_/λ_em_ 485/535 nm. DNA content was correlated to the number of cells through the use of a reference standard curve. hMSCs cultured on flat and topographically-textured microparticle discs were also assessed for metabolic activity using PrestoBlue™ (Invitrogen, UK) at 4 hours post-seeding (for assessment of initial cell attachment; seeded at 3×10^4^ cells/well) and after 1, 5, 8, 12 and 15 days (to assess proliferation; seeded at 1×10^4^ cells/well), according to the manufacturer’s protocol. Briefly, culture medium was replaced by PrestoBlue™:culture medium (1:9) and incubated in the dark for 1.5 hours at 37 °C. Duplicate 100 μL aliquots of supernatant were assessed for fluorescence at at λ_exc_/λ_em_ 560/590 nm. DNA quantification of the lysates collected at days 14 and 21 was carried out using the Quant-iT™ PicoGreen^®^ dsDNA assay (Invitrogen), and concentrations were calculated using a standard curve generated from DNA standard provided. Proliferation was also observed on the discs using immortalized human mesenchymal stem cells (hiMSCs) and imaged using a Nikon SMZ1500 dissection microscope.

### Adhesion Blocking Studies

Cell suspensions in serum-free media were pre-incubated with anti-α_2_ (MAB1950, Merck), anti-β_1_ (MAB1987, Merck), anti-α_5_ (MAB1956Z, Merck), anti-α_v_β_3_ (14-0519, eBioScience) integrin antibodies or anti-mouse isotype IgG antibody (M6898, Sigma-Aldrich) for 30 min at 37°C, prior to seeding in 96-well plates containing discs or control wells pre-coated with 3% (*v/v*) BSA in PBS. Cell suspensions of 4.5×10^5^ cells/mL were used. After incubating for 1 hour at 37°C, non-adherent cells were removed by gently rinsing with PBS several times. The 1 hour period for evaluating cell adhesion was selected to avoid cells secreting a significant amount of matrix nor significantly change their integrin expression profile while adherent(*55*). CyQUANT^®^ NF Cell Proliferation assay was used for cell quantification, expressed as the proportion of cells attached relative to cell numbers attached in the isotype IgG control after subtraction of background values of blank discs.

### Determination of Osteogenesis

For differentiation experiments, 3250 cells were seeded per disc, which corresponds to a range of 2750-10,000 cells/cm^2^.

#### Quantitative Determination of Alkaline Phosphatase Levels

The Alkaline Phosphatase Detection Kit Fluorescence (Sigma-Aldrich, UK) was used according to the manufacturer’s protocol. Cultured cells, 7 days post-seeding, were lysed using 1% Triton X-100 for 20 minutes, followed by three freeze–thaw lysis steps and centrifugation. Samples were incubated with the non-fluorescent 4-methylumbelliferyl phosphate disodium salt as substrate, and resultant fluorescence was measured using a plate reader. ALP activity was normalized to total DNA content, determined using the Quant-iT PicoGreen dsDNA Assay Kit (Invitrogen, UK).

#### Assessment of Osteocalcin Expression

At day 14, hMSCs were rinsed with warm PBS and fixed with 3.7% (w/v) paraformaldehyde in deionized water for 40 min. Cells were permeabilized using 0.1% (w/v) Triton-X 100 in PBS (Sigma-Aldrich) for 30 min. Non-specific binding sites were blocked by incubation in 10% (v/v) normal donkey serum (D9663; Sigma-Aldrich) and 1% (*v/v*) bovine serum albumin (BSA) in PBS for 1 h. Cells were then incubated with anti-Osteocalcin antibody (AB10911; 1:250; Merck Millipore) overnight at 4°C. Donkey anti-rabbit IgG-FITC secondary antibody (1:500; Invitrogen) was added for 2 h. Samples were counterstained with NucBlue^®^ Fixed Cell ReadyProbes (Thermo-Fisher) and visualized using the confocal unit LSM 780 of the Zeiss Elyra PS1 microscope. Quantification of osteocalcin in conditioned media was carried out using the Human Osteocalcin Instant ELISA Kit (eBioscience) at the indicated times according to the manufacturer’s instructions. Data analysis was performed using the elisaanalysis.com software.

#### Assessment of Mineralization

OsteoImage assay (Lonza) was used after 3 weeks of culture. Mineralization was measured quantitatively by spectrophotometer at 492 nm excitation and 520 nm emission wavelengths. Results were normalized to DNA content, quantified using Quant-iT PicoGreen assay (Thermo Fisher Scientific). Presence of extracellular calcium deposits was also verified using von Kossa staining (silver nitrate solution with exposure to UV for 60 min then sodium thiosulfate solution; Sigma Aldrich) and SEM.

#### Gene Expression Array

Total RNA was extracted from hMSCs cultured in basal culture medium for 2 weeks using RNAqueous™-Micro Kit. RNA from each sample was reverse transcribed using SuperScript™ IV First-Strand Synthesis System. RNA quality and quantity were evaluated using Agilent Technologies TapeStation. Quantitative real-time PCR was performed in 96-well TaqMan^®^ Gene Expression Array Cards following manufacturer’s instructions, and run on 7900HT Fast Real-Time PCR System with 384-Well Block Module (ThermoFisher Scientific). Two independent experiments were run (*n*=2), and data was analyzed using DataAssist software v3.01 (Applied Biosystems). Endogenous control normalization was used, with GAPDH and ACTB as selected internal controls. Only genes showing no outlier replicates and a maximum Ct value of 37 were included; *p*-values were adjusted using Benjamini-Hochberg False Discovery Rate (FDR). Genes were reported as differentially expressed if showing a mean fold change of over ±1.5 and FDR <0.1.

#### Scanning Electron Microscopy

Samples were sequentially dehydrated with ethanol, mounted onto aluminum stubs and sputter-coated with gold for 180 s prior to imaging on a JEOL 6060LV variable pressure scanning electron microscope (Jeol UK Ltd.) at 10kV.

#### Microparticle-conditioned media

To prepare microparticle-conditioned media, culture media was incubated with either smooth or dimpled microparticle discs for 14 or 21 days (same length of time used to study OCN expression and mineralization, respectively). This media was periodically and immediately (without storage) transferred to cells cultured on TCP after filtration through a cell strainer (pore size 40 μm), to observe the influence of any residual fusidic acid on differentiation.

#### Extracellular Metabolites Extraction and LC-MS Based Metabolite Profiling

At day 15 post-seeding, 1 mL of culture media were collected and centrifuged. 160 μL were transferred to a new tube for extraction and protein precipitation by addition of 480 μL cold methanol, mixing and incubating at −20 °C for 20 min. Samples were then centrifuged at 16000*xg* at 4°C for 10 min, transferred to pre-cooled tubes and stored at −80 °C. Fresh culture medium samples and methanol samples were processed in parallel as no-cell controls. Samples were prepared as six biological replicates (two donors) and the experiment run in two independent repeats. An equal mixture of all samples was prepared as a pooled quality control for assessment of instrument performance(*56*). For metabolite foot-printing, LC-MS was performed on an Accela LC system coupled to an Exactive MS (ThermoFisher Scientific, UK), as previously described.(*57*) Spectral data was acquired in full scan ion mode (*m/z* 70–1400, resolution 50,000) in positive and negative electrospray ionization modes. Probe temperature and capillary temperature were kept at 150 and 275 °C, respectively. Chromatographic separation was carried out using a ZIC-*p*HILIC (5μm column, 150 mm × 4.6 mm, Merck Sequant) maintained at 45°C and flow rate of 300 μLmin^−1^ as previously described.(*58*) Mobile phase consisted of (A) 20 mM ammonium carbonate in water and (B) 100% acetonitrile, eluted with a linear gradient from 80% B to 5% B over 15 min, followed by a 2 min linear gradient from 5% B to 80% B, and 7 min re-equilibration with 80% B (injection volume= 10 μL; 4°C). LC-MS data were processed using XCMS for untargeted peak-picking(*59*), and peak matching was carried out using mzMatch.(*60*) IDEOM (v20) was used for putative metabolite identification.(*61*) Level 1 metabolite identification was performed by matching accurate masses and retention times of 268 authentic standards, which were analyzed using the same analytical conditions according to the metabolomics standards initiative(*62*, *63*). Level 2 putative identification was considered when standards were unavailable and predicted retention times were used as an orthogonal means to improve metabolite identification. Supervised orthogonal partial least squares discriminant analysis (OPLS-DA) was initially performed using SIMCA-P version 13.0.2 (Umetrics AB, Sweden) for general visualization of metabolite differences and to observe differences between control and treated samples. Quality of the models was evaluated based on *R^2^* (goodness-of-fit) and *Q^2^* (predictive ability). For detection of key discriminatory metabolites, mass ions were selected by variable importance in projection (VIP) values, where VIP >1 were considered potential biomarkers. Univariate analysis was used as a final feature selection. Student *t*-test with false discovery rate (FDR) correction was carried out using MetaboAnalyst(*64*) to evaluate levels of significant differences between controls and microparticle-cultured media, as well as pathway analyses. Analyses were also carried out using Ingenuity Pathway Analysis (IPA; QIAGEN Inc.).(*65*)

#### In vivo Study

Polyimide implant tubes with holes were washed with absolute ethanol in an ultrasound bath for 10 min. After several rinses with ethanol, implant tubes were autoclaved. Autoclaved implant tubes were filled with the microparticles under investigation, using rat-tail collagen type I (First Link Ltd., UK) as carrier. Collagen was freshly prepared by combining stock collagen type I solution (2.05 mg/mL in 0.6% acetic acid) with 3 mL ice-cold 0.1 M sodium hydroxide and 0.5 mL 10x DMEM. Microparticles were sterilized by ultraviolet radiation. Microparticles were prepared in collagen solution at a final concentration of 620 mg/mL, and 3 μL of the microparticle suspensions were placed into the implant tubes. Collagen gel loaded with BMP-2 (75 μg/mL, R&D Systems) was used as a positive control, while plain collagen was used as a negative control. All materials were prepared the day before surgeries.

Experiments were performed under a project license issued under the ASPA (Animals Scientific Procedures Act 1986) by the Home Office, UK. C57B1/6J male mice (8 to 10 weeks old; Charles River laboratories) were anesthetized under isoflurane, and the right forelimb was shaved and swabbed with isopropyl alcohol and iodine solution. After induction of anesthesia, mice were provided with buprenorphine and carprofen administrated subcutaneously. A skin incision was made along the forearm, and muscle tissue over the radius was blunt-dissected. A 2.5-mm defect was created in the center of the radius using a double-bladed bone cutter. Implant tubes were placed into the defect by fitting it at the proximal and distal ends of the radial defect, and the incision was then closed with degradable Vicryl^®^ suture. Mice were monitored after surgery for signs of distress, movement and weight loss. At the end of the experiment (8 weeks), mice were euthanized, and radial bones were explanted and fixed in 10% neutral-buffered formalin solution.

High-resolution micro-X-ray-computed tomography system (micro-CT, Skyscan 1174) was used to determine bone ingrowth within the defects. Fixed samples were scanned at a voltage of 50 kV, current of 800 μA, and a voxel resolution of 9.86 μm. A 0.5mm aluminium filer was also applied. Transmission images were reconstructed using Skyscan supplied software (NRecon) with the resulting two-dimensional image representing a single 9.86 μm slice. Quantitative analysis of bone ingrowth was obtained using direct morphometry calculations in the Skyscan CTAn software package.

Bone samples from all treatment and control groups (*n*=5 each, and two samples with no defect) were decalcified using 5% (*v/v*) formic acid and dehydrated in progressively higher concentrations of ethanol followed by xylene. Samples were then embedded in paraffin wax and sectioned. Sets of eight sections were cut at four levels throughout the implant tube and 200 μm apart, then processed for hematoxylin–eosin, Picrosirius Red and Alcian Blue/Fast Red staining. For hematoxylin–eosin staining, sections were immersed in hematoxylin solution and washed and stained with 10% (*w/v*) eosin solution. For Alcian Blue, slides were submerged in Alcian Blue solution, rinsed in borax and counter-stained with nuclear Fast Red before dehydrating and mounting. Collagen was visualized using the Picrosirius Red Stain Kit (Abcam, ab150681) following the manufacturer’s protocol. All samples were dehydrated with ethanol and xylene solutions and mounted using mounting solution. Representative images of stained sections were obtained using a Nanozoomer (Nanozoomer 2.0 HT, Hamamatsu, Japan) and visualized using the Hamamatsu NanoZoomer Digital Pathology system (NDP.view 2).

#### Statistical Analysis

All values are reported as mean values ± SD. Statistical analyses were performed using GraphPad Prism software (PRISM v8; GraphPad Software, USA). Data was analyzed using non-parametric (Kruskal-Wallis), or parametric one-way/two-way analysis of variance (ANOVA) with Tukey or Dunnett’s *post-hoc* tests, with *p* ≤0.05 considered the threshold for statistical significance. Biological replicates using multiple donors were used in all experiments, as detailed in the figure legends.

## General

We acknowledge Dr. Alison Woodward for assistance with metabolomics, Dr. Robert Markus (University of Nottingham) for assistance with confocal microscopy (equipment funded by Biotechnology and Biological Sciences Research Council [BB/L013827/1]), and Dr. Tim Constantin for assistance with PCR. We also thank the Nanoscale and Microscale Research Centre at University of Nottingham for providing access to SEM and XPS instrumentation.

## Author Contributions

The manuscript was created through contributions of all authors. Mahetab Amer carried out the majority of the experimental work and analysis, collated data and drafted the manuscript. Marta Alvarez-Paino optimized microparticle fabrication, fabricated the majority of microparticles used for experiments and carried out degradation and release studies. Jane McLaren carried out micro-CT reconstructions, contributed to histological preparation of samples and assisted with interpretation of *in vivo* data. Francesco Pappalardo carried out AFM-based characterization and analysis. Sumana Shrestha assisted with microparticle fabrication. Jing Qian Wong assisted with optimizing microparticle fabrication. Sara Trujillo-Munoz carried out the *in vivo* study. Salah AbdelRazig carried out the LC-MS runs for metabolomics analysis. Lee Stevens assisted with BET surface area measurements, and Jong Bong Lee carried out HPLC analysis for the FA release studies. Dong-Hyun Kim provided expertise for the review of metabolomics analysis. Insight into fabrication of textured microparticles was provided by David Needham. Cristina González-García and Manuel Salmeron-Sanchez provided expertise and funding for *in vivo* experiments. The work was conceived, organized and made possible with funding awarded to Morgan Alexander, Kevin Shakesheff, Cameron Alexander and Felicity Rose. Manuscript submission was led by Felicity Rose, Cameron Alexander and Morgan Alexander. All authors have given approval to the final version of the manuscript.

## Data Availability Statement

Datasets generated and/or analyzed during this study are available from the corresponding authors on reasonable request.

## Supplementary Materials

**Supplementary Figure 1.**
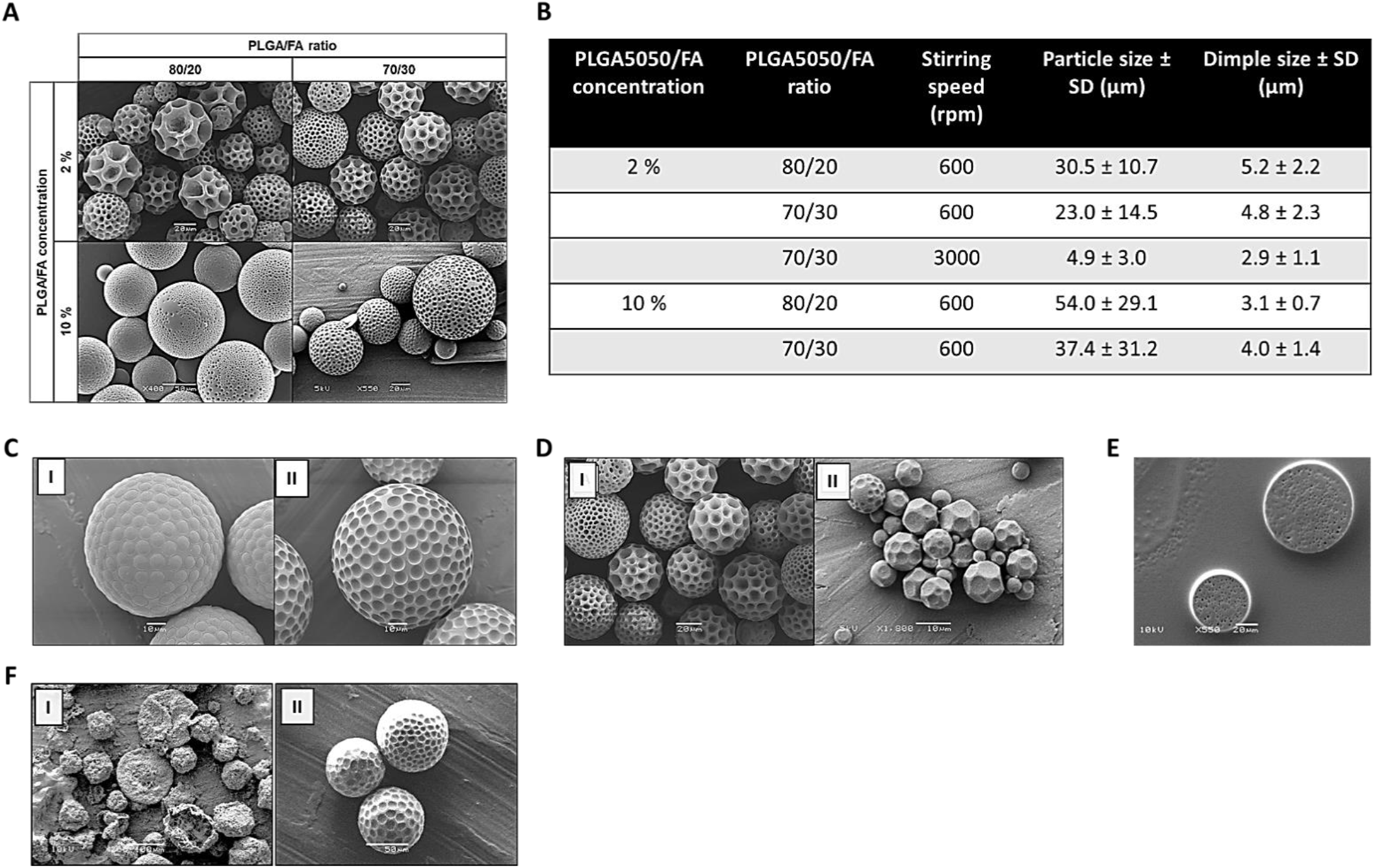
(A) SEM images of dimpled microparticles fabricated at different PLGA5050/ FA ratios and PLGA5050/ FA concentrations in the organic phase (stirring speed 600 rpm). (B) Table displaying average particle size and dimple size of PLGA5050-based microparticles fabricated at different PLGA5050/ FA ratios and PLGA 5050/FA concentrations in the organic phase. (C) SEM images of dimpled microparticles produced at 70/30 PLGA/FA ratios, 10% w/v PLGA5050/FA concentration in the organic phase and 600 rpm stirring speed before (I) and after (II) FA release (Scale bar: 10 μm). (D) SEM images of dimpled microparticles fabricated at 70/30 PLGA/FA ratio, 2% PLGA5050/ FA concentration in the organic phase and a stirring speed of 600 rpm (I) or 3000 rpm (II). (E) Cross-sectional SEM image of dimpled PLGA5050/FA microparticles fabricated at 10% PLGA5050/FA concentration in the organic phase, 80/20 PLGA5050/ FA ratio and stirring speed 600 rpm showing a largely non-porous internal structure with a few sub-micron pores (Scale bar: 20 μm). (F) Degradation studies of PLGA5050/ FA (I) and PLA/ FA (II) microparticles (containing 30% FA), maintained in cell culture DMEM media at 37 °C for 2 months (Scale bar: 100 and 50 μm, respectively).

**Supplementary Figure 2.**
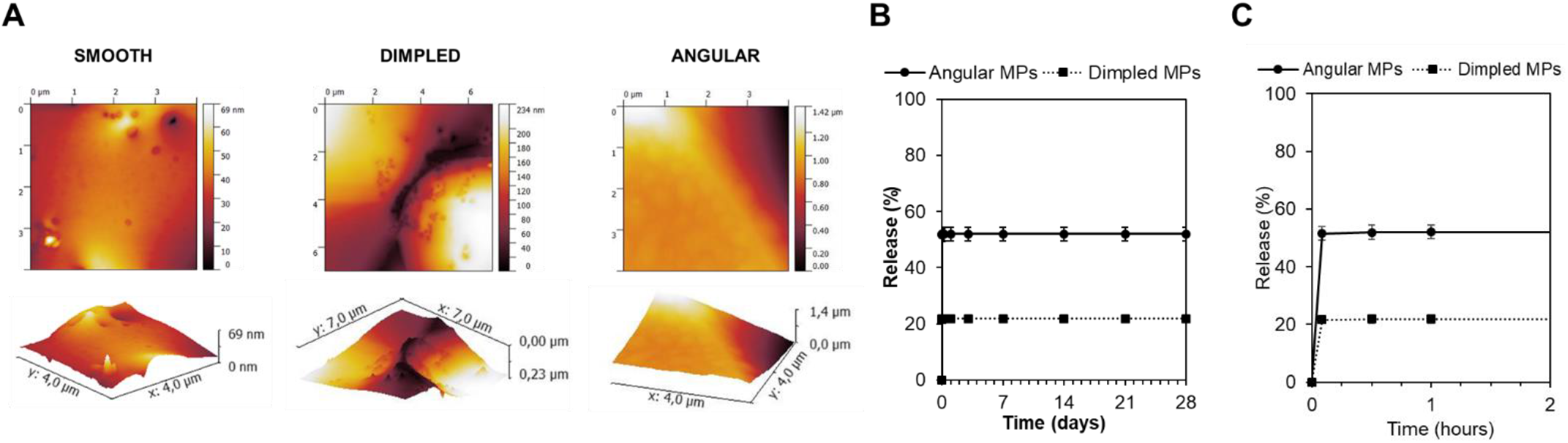
(A) AFM topography images of the different microparticle designs, displaying the scale bars below each image. (B-C) Cumulative release profile of FA in PBS at 37 °C of PLA dimpled (square) and angled (circle) microparticles (n = 4) over 28 days (B) and over the first 2 hours (C).

**Supplementary Figure 3.**
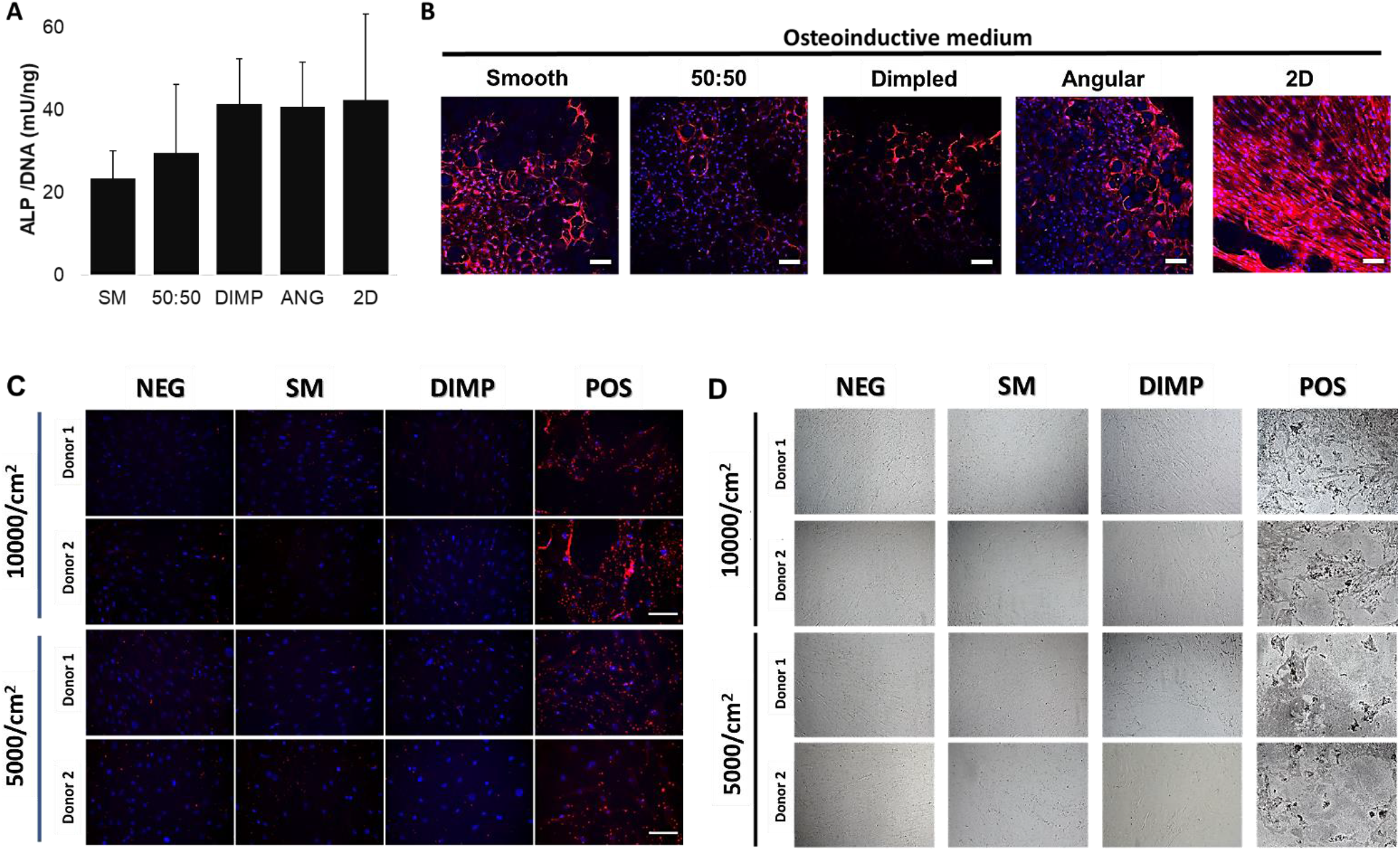
(A) Secreted ALP activity levels in culture media, after normalizing to DNA content, of primary hMSCs cultured in basal media with no osteogenic supplements for 7 days. Values shown are mean ± SD (*n* = 4 in two donors). (B) Maximum intensity projection images of *z*-stack sections obtained by immunostaining and confocal microscopy of primary hMSCs cultured on the various ‘2.5D’ discs in osteoinductive media for 14 days. Cell nuclei (Hoechst) are shown in blue and osteocalcin (OCN) in red. Microparticles, especially textured ones, were slightly auto-fluorescent in the blue channel (Scale bar = 100 μm). (C) Fluorescence microscopy images of primary hMSCs cultured using smooth and dimpled microparticle-conditioned media (two cell seeding densities) on TCP for 14 days. (D) Bright-field microscopy images of von Kossa staining carried out on primary hMSCs cultured using smooth and dimpled microparticle-conditioned media (two cell seeding densities) on TCP for 21 days. Negative and positive controls are included for reference. SM: Smooth microparticle discs; DIMP: Dimpled microparticle discs; 50:50: Equal mixture of smooth and dimpled microparticles in discs; ANG: Angular microparticle discs; 2D: melted microparticle discs.

**Supplementary Figure 4.**
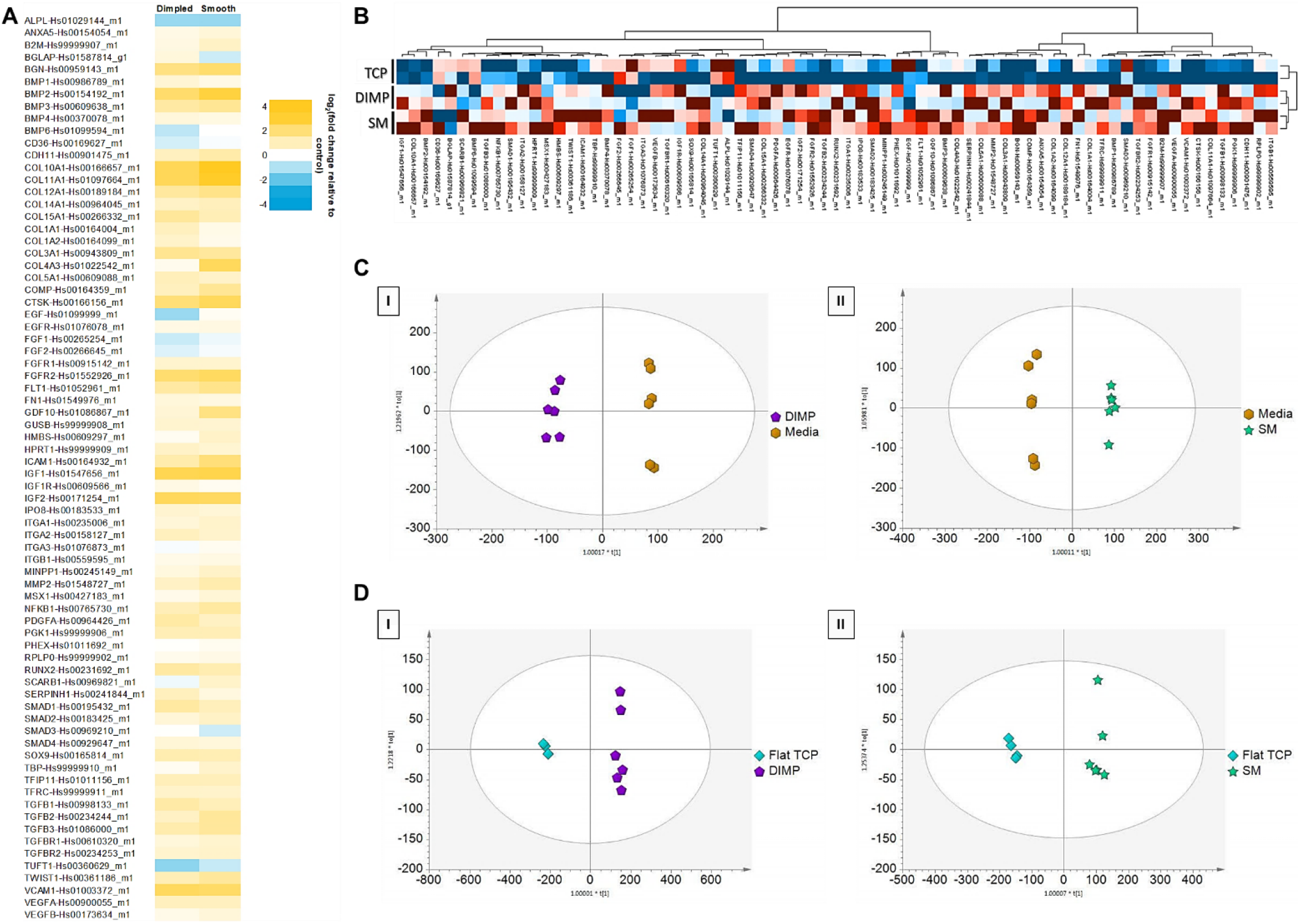
(A) A full list of genes profiled, displaying up- and down-regulated genes in hMSCs cultured on smooth and dimpled microparticles relative to control (cultured on TCP). Relative expression for each gene are plotted as log2 (fold change) relative control after normalization against housekeeping genes. Yellow indicates upregulated and blue indicates downregulated genes (*n* = 2; 2 donors; 2 independent experiments). (B) Hierarchical clustering of gene expression in hMSCs cultured on smooth and dimpled microparticles as well as 2D TCP controls using DataAssist software (*n* = 2 each; Euclidean distance, complete linkage). (C) OPLS-DA scores plots based on metabolic profiling of culture media of dimpled (DIMP; I; R^2^=0.99, Q^2^=0.93) and smooth (SM; II; R^2^=0.996, Q^2^=0.97) microparticles versus fresh media samples two weeks post-seeding. (D) OPLS-DA scores plots based on metabolic profiling of culture media of dimpled (DIMP; I; R^2^Y=0.996, Q^2^=0.99) and smooth (SM; II; R^2^=0.99, Q^2^=0.98) microparticles versus flat TCP-cultured samples two weeks post-seeding.

**Supplementary Figure 5.**
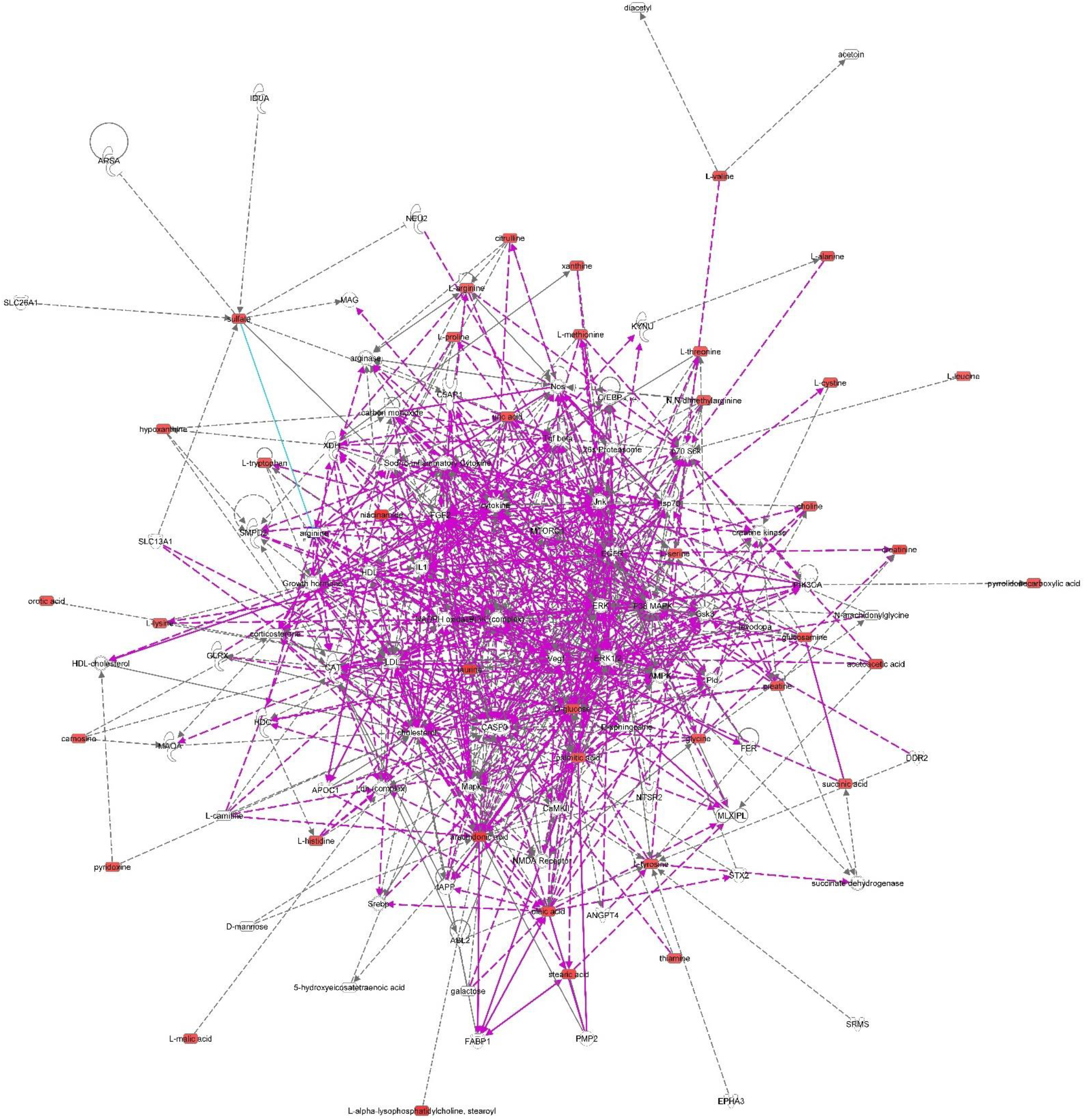
Merged network combining major signalling networks associated with the differentially expressed metabolites in smooth versus dimpled microparticle samples, as identified via ingenuity pathway analysis (*n*=6 each; 2 independent experiments using two donors).

**Supplementary Figure 6.**
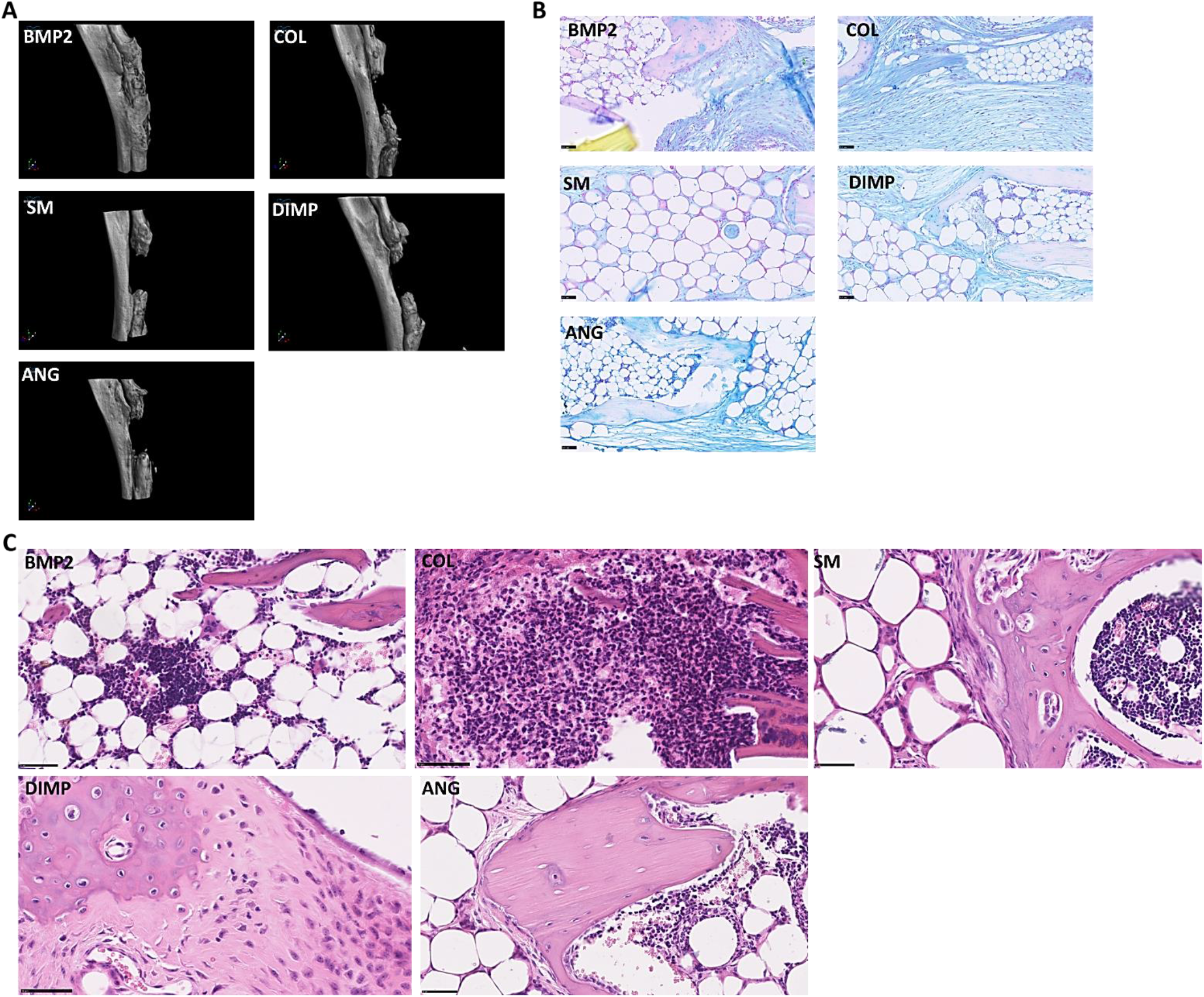
(A) Three-dimensional μCT reconstructions after 8 weeks, with three microparticle designs and two controls (B) Representative histological images showing sections of 8-week radial samples stained with Alcian Blue/Fast Red (Scale bar: 50 μm) to detect the presence and distribution of sulphated glycosaminoglycans. (C) Further magnification of the areas of interest within the sections of 8-week radial samples stained with H&E displayed in Figure 6 (scale bar: 50 μm). SM: smooth, DIMP: dimpled, ANG: angular, BMP2 as positive control, COL: collagen-only control.

**Supplementary Table 1.** Morphological characterization of dimpled microparticles fabricated by varying emulsion method parameters (microparticle properties shown are mean values ± SD)

| EMULSION SETTINGS |  |  | MICROPARTICLE PROPERTIES |  |
| --- | --- | --- | --- | --- |
| Stirring speed (RPM) | Polymer concentration (%) | PLA/FA ratio | Particle size ± SD (µm) | Dimple size ± SD (µm) |
| 1600 | 15 | 90/10 | 48.4 ± 18.9 | 2.56 ± 1.0 |
| 1600 | 15 | 75/25 | 46.9 ± 17.5 | 7.6 ± 2.8 |
| 1600 | 15 | 60/40 | 43.8 ± 20.4 | 11.9 ± 5.1 |
| 1600 | 15 | 40/60 | 40.8 ± 14.3 | 14.6 ± 5.1 |
| 600 | 10 | 80/20 | 61.7 ± 23.0 | 5.3 ± 2.6 |
| 600 | 10 | 70/30 | 58.5 ± 13.2 | 5.2 ± 2.2 |
| 600 | 10 | 60/40 | 72.8 ± 12.7 | 7.5 ± 3.5 |

**Supplementary Table 2.** Calculation of the cell-seeded surface areas of various microparticle-containing discs, assuming monolayer coverage and monodispersed size distribution.

|  | SMOOTH | DIMPLED | ANGULAR |
| --- | --- | --- | --- |
| BET SA (m<sup>2</sup>/g) | 0.0486 | 0.1002 | 0.0944 |
| Diameter (µm) | 58.2 | 58.5 | 43.4 |
| Number of microparticles/mg<br>*(calculated using the equation*<br>*$N = 6 \times 10^{12} / (\pi \cdot \rho_s \cdot d^3) )$ <sup>#</sup>* | 7750 | 7632 | 18691 |
| Circle packing calculations<br>(microparticles/disc) | 10158 | 10054 | 18329 |
| Mass of microparticles/disc<br>(mg) | 1.31 | 1.32 | 0.98 |
| Surface area of particles/disc (cm<sup>2</sup>) | 0.64 | 1.32 | 0.93 |
| Area of disc's surface (on which cells are seeded; cm<sup>2</sup>) | 0.32 | 0.66 | 0.47 |

<sup>#</sup> N= number of microparticles/gram; ρ<sub>s</sub> = density of PLA (1.25 g/cm<sup>3</sup>); d = mean diameter (µm)

**Supplementary Table 3.** Identification of differential metabolites detected using LC/MS data with ≥1.5 relative intensity of dimpled samples relative to smooth and normalised to DNA content, presented as relative intensity to smooth microparticle samples (*n*= 6 of each microparticle type and *n*=4 TCP-cultured samples).

| Metabolite | Chemical formula | Pathway | Class | MSI Level<sup>#</sup> | DIMP | TCP |
| --- | --- | --- | --- | --- | --- | --- |
|  |  |  |  |  | Relative Intensity vs SM | Relative Intensity vs SM |
| (5Z,7E)-<br>(1S,3R,20R,22R)-<br>24a,24b-dihomo-9,10-seco-5,7,10(19)-<br>cholestatrien-23-yne-1,3,22,25-tetrol | C<sub>29</sub>H<sub>44</sub>O<sub>4</sub> | Lipids: Sterol lipids | Secosteroids | 2 | 194.8 | 0.09 |
| PG(20:1(11Z)/0:0) | C<sub>26</sub>H<sub>51</sub>O<sub>9</sub>P | Lipids:<br>Glycerophospholipids | Glycerophosphoglycerols | 2 | 70.6 | 0 |
| Nicotinamide | C<sub>6</sub>H<sub>6</sub>N<sub>2</sub>O | Metabolism of Cofactors<br>and Vitamins | Nicotinate & nicotinamide<br>metabolism | 1 | 1.79 | 0.41 |
| LysoPC(18:0) | C<sub>26</sub>H<sub>54</sub>NO<sub>7</sub>P | Lipids:<br>Glycerophospholipids | Glycerophosphocholines | 2 | 1.76 | 0.33 |
| Arachidonic acid | C<sub>20</sub>H<sub>32</sub>O<sub>2</sub> | Lipids: Fatty Acyls | Fatty Acids & Conjugates | 2 | 1.73 | 0.27 |
| Taurine | C<sub>2</sub>H<sub>7</sub>NO<sub>3</sub>S | Lipid Metabolism | Taurine & hypotaurine<br>metabolism | 1 | 1.71 | 0.35 |
| 5-Oxo-D-proline | C<sub>5</sub>H<sub>7</sub>NO<sub>3</sub> | Amino Acid Metabolism | D-Glutamine & D-glutamate metabolism | 2 | 1.68 | 0.43 |
| Butylamine | C<sub>4</sub>H<sub>11</sub>N | - | - | 2 | 1.66 | 0.46 |
| 6R-(8-hydroxydecyl)-<br>2R-(hydroxymethyl)-<br>piperidin-3R-ol | C<sub>16</sub>H<sub>33</sub>NO<sub>3</sub> | Lipids: Sphingolipids | Sphingoid bases | 2 | 1.65 | 0.43 |
| L-3-Amino-<br>isobutanoate | C<sub>4</sub>H<sub>9</sub>NO<sub>2</sub> | Amino Acid Metabolism | Valine, leucine &<br>isoleucine degradation | 2 | 1.65 | 0.40 |
| 2-Amino-tetradecanoic<br>acid | C<sub>14</sub>H<sub>29</sub>NO<sub>2</sub> | Lipids: Fatty Acyls | Amino Fatty Acids | 2 | 1.64 | 0.44 |
| L-Thiazolidine-4-carboxylate | C<sub>4</sub>H<sub>7</sub>NO<sub>2</sub>S | - | - | 2 | 1.62 | 0.87 |
| **2-Hydroxy-2,4-pentadienoate** | C<sub>5</sub>H<sub>6</sub>O<sub>3</sub> | Amino Acid Metabolism | Phenylalanine metabolism | 2 | 1.61 | 0.19 |
| **Stearic acid** | C<sub>18</sub>H<sub>36</sub>O<sub>2</sub> | Lipid Metabolism | Fatty acid biosynthesis | 1 | 1.60 | 0.44 |
| **Acetoacetate** | C<sub>4</sub>H<sub>6</sub>O<sub>3</sub> | Lipid Metabolism | Synthesis & degradation of ketone bodies; Valine, leucine & isoleucine degradation; Tyrosine metabolism | 2 | 1.59 | 0.46 |
| **2R-Amino-butanoic acid** | C<sub>4</sub>H<sub>9</sub>NO<sub>2</sub> | Lipids: Fatty Acyls | Amino Fatty Acids | 2 | 1.59 | 0.43 |
| **Oleic acid** | C<sub>18</sub>H<sub>34</sub>O<sub>2</sub> | Lipids: Fatty Acyls | Fatty acid biosynthesis | 2 | 1.58 | 0.42 |
| **3-Ethylmalate** | C<sub>6</sub>H<sub>10</sub>O<sub>5</sub> | Carbohydrate Metabolism | Glyoxylate & dicarboxylate metabolism | 2 | 1.58 | 0.01 |
| **4-Oxo-4-(3-pyridyl)-butanoate** | C<sub>9</sub>H<sub>9</sub>NO<sub>3</sub> | - | Nicotine degradation | 2 | 1.58 | 0.46 |
| **Creatinine** | C<sub>4</sub>H<sub>7</sub>N<sub>3</sub>O | Amino Acid Metabolism | Arginine and proline metabolism | 1 | 1.58 | 0.41 |
| **2-Methylpropanamine** | C<sub>4</sub>H<sub>11</sub>N | - | - | 2 | 1.58 | 0.42 |
| **D-Fructose** | C<sub>6</sub>H<sub>12</sub>O<sub>6</sub> | Carbohydrate Metabolism | Fructose and mannose metabolism; Galactose metabolism | 1 | 1.56 | 0.38 |
| **L-1-Pyrroline-3-hydroxy-5-carboxylate** | C<sub>5</sub>H<sub>7</sub>NO<sub>3</sub> | Amino Acid Metabolism | Arginine and proline metabolism | 2 | 1.56 | 0.42 |
| **N-(4-nitrophenyl)validamine** | C<sub>13</sub>H<sub>18</sub>N<sub>2</sub>O<sub>6</sub> | - | - | 2 | 1.56 | 0.42 |
| **D-Glucose** | C<sub>6</sub>H<sub>12</sub>O<sub>6</sub> | Carbohydrate Metabolism | Glycolysis/ Gluconeogenesis; Pentose phosphate pathway; Galactose metabolism | 1 | 1.55 | 0.18 |
| **D-Glucosamine** | C<sub>6</sub>H<sub>13</sub>NO<sub>5</sub> | Carbohydrate Metabolism | Amino sugars metabolism | 1 | 1.55 | 0.35 |
| **Methyl5-(but-3-en-1-yl)amino-1,3,4-oxadiazole-2-carboxylate** | C<sub>8</sub>H<sub>11</sub>N<sub>3</sub>O<sub>3</sub> | - | - | 2 | 1.55 | 0.34 |
| **Pyridoxine** | C<sub>8</sub>H<sub>11</sub>NO<sub>3</sub> | Metabolism of Cofactors & Vitamins | Vitamin B6 metabolism | 1 | 1.54 | 0.36 |
| **L-Alanine** | C<sub>3</sub>H<sub>7</sub>NO<sub>2</sub> | Amino Acid Metabolism | Alanine & aspartate metabolism; Cysteine metabolism; Taurine and hypotaurine metabolism | 1 | 1.54 | 0.59 |
| **4-Acetamidobutanoate** | C<sub>6</sub>H<sub>11</sub>NO<sub>3</sub> | Amino Acid Metabolism | Arginine and proline metabolism | 2 | 1.53 | 0.43 |
| **2,3-Dimethylmaleate** | C<sub>6</sub>H<sub>8</sub>O<sub>4</sub> | Metabolism of Cofactors & Vitamins | Nicotinate & nicotinamide metabolism | 2 | 1.53 | 0.36 |
| **2-C-Methyl-D-erythritol 4-phosphate** | C<sub>5</sub>H<sub>13</sub>O<sub>7</sub>P | Lipid Metabolism | Biosynthesis of steroids | 2 | 1.53 | 0.39 |
| **4-(3-pyridyl)-butanoate** | C<sub>9</sub>H<sub>11</sub>NO<sub>2</sub> | - | Nicotine degradation | 2 | 1.53 | 0.40 |
| **N,N-Dimethylglycine** | C<sub>4</sub>H<sub>9</sub>NO<sub>2</sub> | Amino Acid Metabolism | Glycine, serine & threonine metabolism | 2 | 1.52 | 0.40 |
| **Sulfate** | H<sub>2</sub>O<sub>4</sub>S | Nucleotide Metabolism | Purine metabolism; Cysteine metabolism; Sulfur metabolism | 2 | 1.52 | 0.40 |
| **N-Acetyl-L-histidine** | C<sub>8</sub>H<sub>11</sub>N<sub>3</sub>O<sub>3</sub> | - | - | 2 | 1.52 | 0.40 |
| **(S)-Malate** | C<sub>4</sub>H<sub>6</sub>O<sub>5</sub> | Carbohydrate Metabolism | TCA cycle; Glutamate metabolism; Alanine and aspartate metabolism; Pyruvate metabolism | 1 | 1.51 | 0.43 |
| **Urate** | C<sub>5</sub>H<sub>4</sub>N<sub>4</sub>O<sub>3</sub> | Nucleotide metabolism | Purine metabolism | 1 | 1.50 | 0.40 |
| **Hexadecanoic acid** | C<sub>16</sub>H<sub>32</sub>O<sub>2</sub> | Lipid metabolism | Fatty acid biosynthesis; Fatty acid elongation in mitochondria; Fatty acid metabolism; Biosynthesis of unsaturated fatty acids | 2 | 1.50 | 0.38 |
| **L-Citrulline** | C<sub>6</sub>H<sub>13</sub>N<sub>3</sub>O<sub>3</sub> | Amino acid metabolism | Arginine and proline metabolism | 1 | 1.50 | 0.35 |
| **Creatine** | C<sub>4</sub>H<sub>9</sub>N<sub>3</sub>O<sub>2</sub> | Amino acid metabolism | Glycine, serine and threonine metabolism; Arginine and proline metabolism | 1 | 1.50 | 0.43 |

<sup>#</sup> Level 1 confidence (according to the Metabolomics Standards Initiative [MSI]) was based on accurate mass and retention times for metabolites. Other metabolites were putatively annotated (MSI level 2) based on accurate mass and predicted retention time using IDEOM database (only IDEOM confidence scores ≥5 were included).

